# Value-guided remapping of sensory circuits by lateral orbitofrontal cortex in reversal learning

**DOI:** 10.1101/2020.03.11.982744

**Authors:** Abhishek Banerjee, Giuseppe Parente, Jasper Teutsch, Christopher Lewis, Fabian F. Voigt, Fritjof Helmchen

**Affiliations:** Laboratory of Neural Circuit Dynamics, Brain Research Institute, University of Zürich; Neuroscience Center Zürich, Winterthurerstrasse 190, 8057 Zürich, Switzerland; Biosciences Institute, Newcastle University, United Kingdom

**Keywords:** Orbitofrontal cortex, Sensory processing, Reversal learning, S1, Reward-history, Credit assignment, Rule-switch

## Abstract

Flexible decision-making is crucial for adaptive behaviour. Such behaviour in mammals largely relies on the frontal cortex, and specifically, the orbitofrontal cortex (OFC). How OFC neurons encode decision variables and instruct sensory areas to guide adaptive behaviour is a key open question. Here we developed a reversal learning task for head-fixed mice together with two-photon calcium imaging to monitor the activity of lateral OFC neuronal populations and investigated their dynamic interaction with primary somatosensory cortex (S1). Mice trained on this task learned to discriminate go/no-go tactile stimuli and adapt their behaviour upon changes in stimulus–reward contingencies (‘rule-switch’). Longitudinal imaging at cellular resolution across weeks during all behavioural phases revealed a distinct engagement of S1 and lateral OFC neurons: S1 neural activity reflected task learning-related responses, while neurons in the lateral OFC saliently and transiently responded to the rule-switch. A subset of OFC neurons conveyed a value prediction error signal via feedback projections to S1, as direct anatomical long-range projections were revealed by retrograde tracing combined with whole-brain light-sheet microscopy. Top-down signals implemented an update of sensory representations and functionally reconfigured a small subpopulation of S1 neurons that were differentially modulated by reward-history. Functional remapping of these neurons crucially depended on top-down inputs, as chemogenetic silencing of lateral OFC neurons disrupted reversal learning and impaired plastic changes in these outcome-sensitive S1 neurons. Our results reveal the presence of long-range cortical interactions between cellular ensembles in higher and lower-order brain areas specifically recruited during context-dependent learning and task-switching. Such interactions crucially implement history-dependent reward-value computations and error heuristics, which, in turn, help guide adaptive behaviour.

## Main Text

Animals flexibly adapt their behaviours in response to variable contextual changes in the environment. One prototypical adaptive behaviour is value-guided decision-making, which relies on diverse neural systems thought to realise key computations, such as the assignment of value to specific stimulus-action pairs based on reward-history. Failure to adapt behaviour is a common symptom in many neurological disorders such as autism and schizophrenia^1^. In mammals, value-guided decision-making is governed by distributed neural circuits engaging cognitive primitives in prefrontal areas of the neocortex^2,3^. The orbitofrontal cortex (OFC) is particularly implicated in cognitive evaluation of stimulus-outcome associations^4–7^. OFC is a key higher-order association area that has been shown to have extensive connections with sensory cortices as well as subcortical structures of the reward system^8,9^. However, how neuronal populations in OFC encode decision-related variables and dynamically engage in flexible decision-making upon changes in reward contingencies is poorly understood. It is also unclear whether and how OFC neurons hierarchically instruct sensory areas to remap their activity for the refinement of sensory representations to support value-guided adaptive behaviour.

To study the neural dynamics underlying flexible decision-making, we employed a reversal learning paradigm based on tactile discrimination. We trained mice in a whisker-based ‘go/no-go’ texture-discrimination task^10^ (**Fig. 1a**; coarse P100 sandpaper as go-texture vs. smooth P1200 sandpaper as no-go-texture; **Methods**). Once task performance reached expert level (discriminability index *(d’)* > 1.5; **Methods**), we implemented a ‘rule-switch’ by reversing stimulus-reward contingency (**Fig. 1b**). Mice reached high *d’* values during initial learning (from ‘learning naïve’, *LN*, through ‘learning expert’, *LE*), decreased performance after reversal, and regained high *d’* values after re-learning the task (from ‘reversal naïve’, *RN*, through ‘reversal expert’, *RE*) (**Fig. 1c, Supplementary Fig. 1**, n = 11 mice). Reversal learning was significantly faster than the initial learning and the performance remained stable over weeks (**Fig. 1d, Supplementary Fig. 1**). Task performance depended on sensory input and was independent of initial go-texture (n = 3 mice trained first on P1200 texture; **Supplementary Fig. 1**). Mice developed anticipatory whisking as well as well-timed licking during initial learning^11^. Following the rule-switch, the overall whisking kinematics did not change as they maintained the whisking-for-touch strategy but transiently reverted to uncertain and delayed licking before regaining confidence in the *RE* period (**Supplementary Fig. 2**).

**Figure 1.**
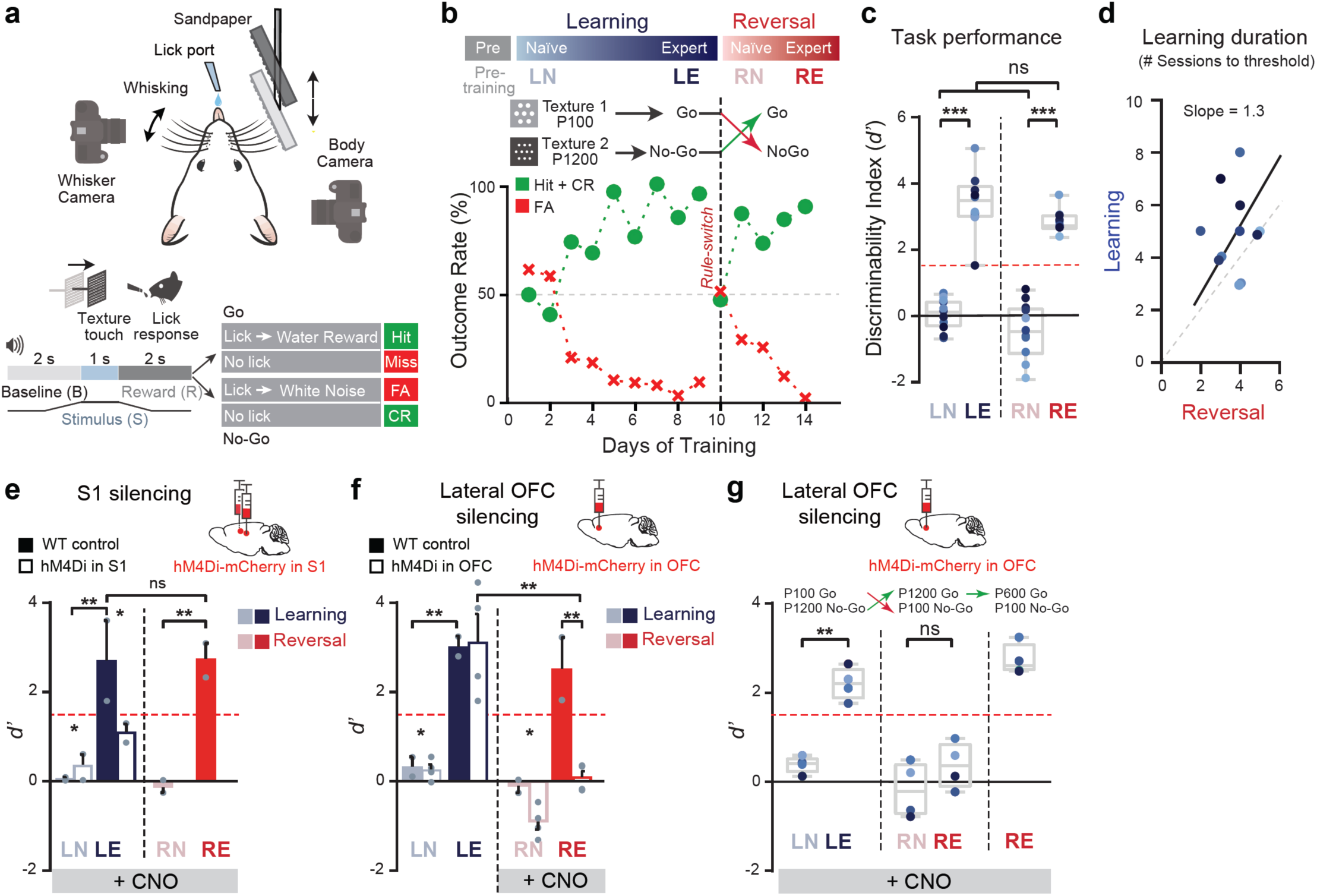
Lateral OFC-dependent reversal learning in a texture-discrimination task. **a**, Top: A schematic diagram of experimental setup. Bottom: Trial structure and types of trial outcome (CR, correct rejection; FA, false alarm). **b**, Time course of task performance of an example mouse measured as mean correct rate (Hit + CR) and FA rate. After reaching stable high performance, stimulus-reward contingency was reversed (‘rule-switch’). Top: Definition of salient task periods (*LN*: learning naïve, *LE*: learning expert, *RN*: reversal naïve, *RE*: reversal expert). **c**, Performance (*d’* values) in the four task periods pooled across 11 mice (different blue shadings), all of which completed reversal learning. Box plots showing median, 25^th^ and 75^th^ percentiles as box edges, and minimum and maximum as whiskers. **d**, Number of sessions to reach expert threshold (*d’* > 1.5) for initial learning plotted against reversal learning. **e**, Virus-induced expression of inhibitory DREADD (hM4Di) in S1 (n = 3 mice). Silencing S1 by daily systemic CNO applications prevented mice from reaching expert threshold (*d’* < 1.5 in *LE*; hence these mice were not reversed). CNO-treated wild-type control mice not expressing hM4Di (WT, n = 4) showed normal learning and reversal learning. **f**, Virus-induced expression of hM4Di in lateral OFC (n = 4 mice). Silencing of lateral OFC during *RN* and *RE* periods impaired reversal learning. **g**, Silencing lateral OFC throughout all task phases did not affect initial discrimination learning but impaired reversal learning after rule-switch (n = 4 mice). Mice were, however, still able to learn a new stimulus-outcome association (novel P600 sandpaper as go-texture). All data presented as mean ± S.E.M., *p < 0.05, **p < 0.01, Wilcoxon rank-sum test.

We focused on two cortical areas implicated in task-learning: barrel cortex in the primary somatosensory cortex (S1), which is important for tactile discrimination and sensory learning^12^, and the lateral OFC, for its role in credit-assignment^8^. To examine the behavioural and causal relevance of these two areas, we expressed inhibitory DREADD receptors (hM4Di) in excitatory neurons in either S1 or lateral OFC (for histology and electrophysiological validation, see **Supplementary Figs. 3** and **4**). Inhibition of S1 neurons during initial training (via daily CNO injections before each behavioural training sessions during *LN* and *LE* periods) prevented mice from learning the discrimination task (**Fig. 1f**). On the other hand, inhibiting neurons in the lateral OFC, but not medial OFC^7^, after the rule-switch (during *RN* and *RE* periods) impaired reversal learning and increased perseverative errors (**Fig. 1f-g, Supplementary Fig. 3**). Interestingly, lateral OFC-silenced mice were still able to learn a new stimulus-outcome association after introduction of a novel P600 sandpaper as rewarded go-texture (**Fig. 1g**). Overall, these results indicate a dissociation of neural mechanisms of learning and reversal learning distributed across cortical areas involving the S1 and lateral OFC, respectively.

To directly probe neuronal activity in lateral OFC and S1 during learning and reversal learning, we performed *in vivo* two-photon Ca^2+^ imaging in transgenic mice expressing GCaMP6f in superficial excitatory layer (L)2/3 neurons. We targeted lateral OFC, which resides deep in the frontal cortex^13,14^, by imaging via a gradient-index lens that was inserted through a chronically implanted cannula (**Fig. 2a**; **Supplementary Fig. 5**; **Methods**, n = 4 mice). Cannula-implanted mice showed no noticeable impairments in whisking and in simple behavioural tests (**Supplementary Fig. 5**). We observed large Ca^2+^ transients in lateral OFC neurons with trial-related activity especially during the reward-outcome window (**Fig. 2a**). An example neuron that was longitudinally measured across all behavioural phases displayed large and robust Ca^2+^ responses to the unexpected rewards for the new go-texture after the rule-switch (*RN*), whereas only a modest increase in reward-related activity was found during initial learning (*LE*) (**Fig. 2b**). This high *RN* activity was transient and decreased as the task performance increased during the *RE* period. The same response pattern was evident when averaging across all OFC neurons, with a significant increase in reward-related Ca^2+^ transient amplitude after the rule-switch (*LE*→*RN;* **Fig. 2c**). These findings are consistent with lateral OFC developing a representation of outcome-value that is strongly visible upon rule-switch^15^. In line with this notion, OFC neuronal responses to a third texture type (P600), associated with a constant small reward both before and after reversal, remained unchanged in control experiments (**Supplementary Fig. 6**). In stark contrast, L2/3 neurons in S1, imaged through a chronic cranial window (n = 5 mice), exhibited abundant Ca^2+^ activity during both the stimulus-presentation as well as the reward-outcome window (**Fig. 2d**). Rewarded go-texture-related Ca^2+^ transients emerged during initial task-learning (*LN*→*LE*), decreased directly after the rule-switch, and re-appeared for the new go-texture in the *RE* period (an example neuron in **Fig. 2e**). This response remapping was also observed when averaging across all S1 L2/3 neurons (**Fig. 2f**). Notably, similar response profile was also found in anatomically identified S1→OFC projection neurons targeted via a dual viral labelling strategy (n = 3 mice; **Supplementary Fig. 7**). Together with neural responses, the fraction of active neurons was highest for *LE* and *RN* phases in the lateral OFC and *LE* and *RE* phases in S1, suggesting strong involvement and dissociation of the two regions in these respective salient behavioural phases (**Fig. 2c** and **f**).

**Figure 2.**
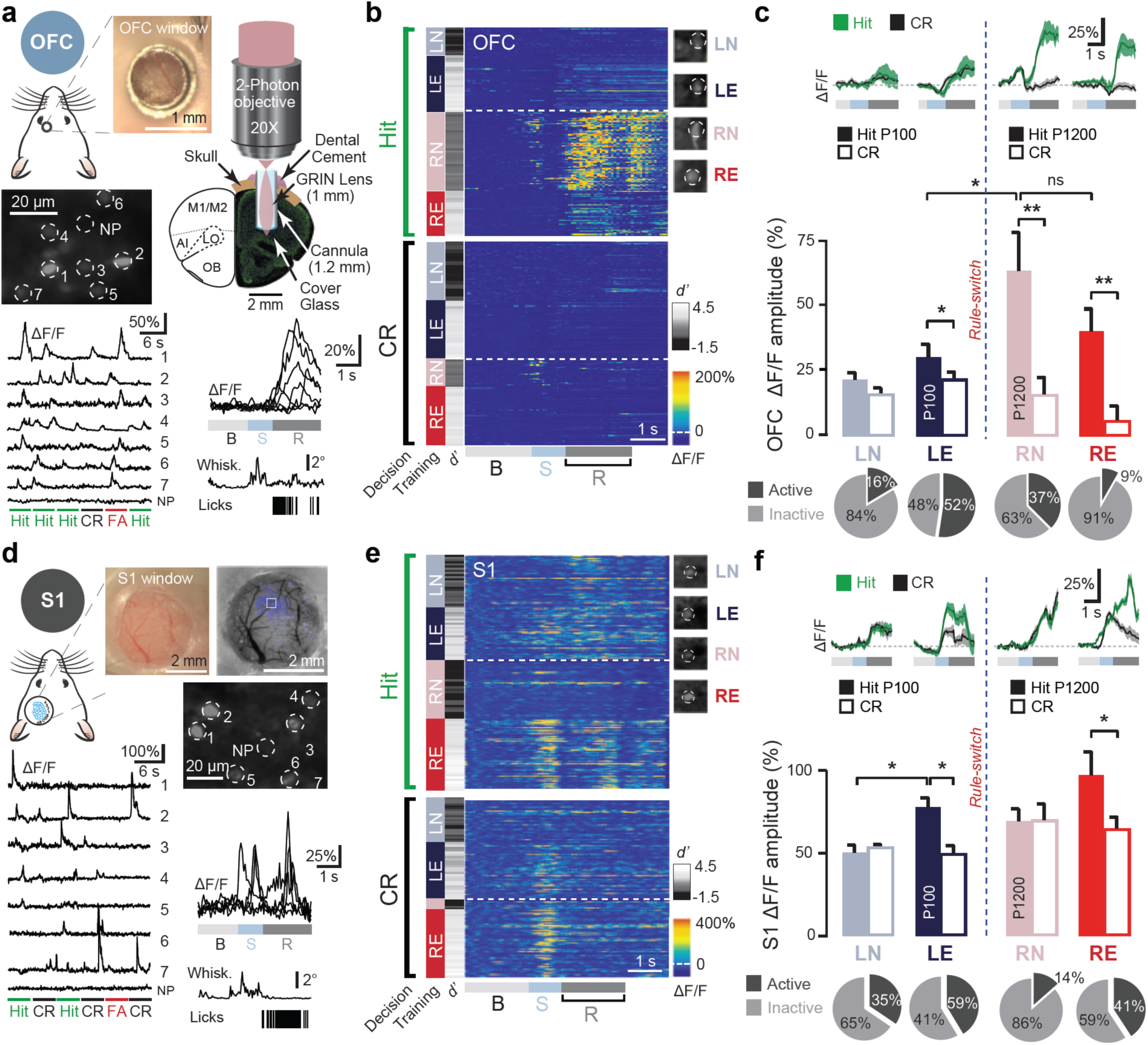
*In vivo* Ca^2+^ imaging of lateral OFC and S1 neurons during reversal learning. **a**, Top: A schematic diagram and picture of cannula-window for imaging in OFC. Bottom: Two-photon fluorescence image and GCaMP6f signals (ΔF/F) during different trial types for example OFC L2/3 neurons imaged through a GRIN lens. Lower right: Example Ca^2+^ transients during hit trials for an individual OFC neuron. Example of single-trial time course of whisking-amplitude and lick events during a hit-trial displayed below. B: baseline, S: stimulus-presentation window, R: reward-outcome window. **b**, Heat-map of single-trial ΔF/F responses (sorted by hit and CR; FA and misses not shown) of an example lateral OFC neuron chronically imaged across behavioural phases. Animal performance (*d’*) indicated next to behavioural phases. **c**, Average Ca^2+^ transient amplitude during the reward-outcome window for lateral OFC neurons for hit and CR trials (n = 63 active neurons out of 228 neurons recorded in three mice; n = 15 sessions). Across-trial average Ca^2+^ transients and percentage of active neurons for each behavioural period are shown above and below. **d**, Top: Schematic and photograph of cranial window above S1. Barrel cortex was identified by intrinsic optical signal mapping of whisker-evoked responses (field of view for two-photon imaging indicated). Bottom: Fluorescence image and GCaMP6f signals (ΔF/F) during different trial types for example S1 L2/3 neurons. Bottom right: Example Ca^2+^ transients during hit trials for an example S1 neuron, exhibiting responses during both stimulus-presentation and reward-outcome window, and single-trial time course of whisking-amplitude and lick events. **e**, Heat-map of ΔF/F transients for an example S1 L2/3 neuron as in (b). **f**, Average Ca^2+^ transient amplitude in the reward-outcome window for S1 neurons for hit and CR trials (n = 261 active neurons out of n = 539 recorded neurons in 5 mice; n = 56 sessions; 11 sessions discarded due to motion artefacts). S1 neurons show a selective increase towards hit trials during both expert phases (*LE* and *RE*). Across-trial average Ca^2+^ transients and percentage of active neurons for each behavioural period are shown above and below. All data presented as mean ± S.E.M.; *p < 0.05, **p < 0.01 Wilcoxon rank-sum test.

Taking advantage of longitudinal activity measurements from the same neurons across weeks, we further examined the heterogeneity of neuronal responses in lateral OFC and S1 and their stability or flexibility upon rule-switch. A key question is whether neurons which preferentially respond to hit-trials after initial learning will retain response selectivity for the new or the old go-texture after reversal, i.e. whether they remain more stimulus-selective or outcome-selective. To quantify response selectivity of active neurons, we defined an ROC-based hit/CR selectivity index (*SI*) that ranges from -1 and +1 (significance tested by permutation test, p < 0.05; **Methods**; **Supplementary Fig. 8**)^16^. We focused on hit/CR *SI* values for the reward-outcome window. Note that the hit/CR *SI per se* cannot distinguish between stimulus- and outcome-selectivity because these trial-types differ in both texture-type and action-outcome. However, by comparing *SI* values before and after the rule-switch we can reveal whether a neuron’s new-hit/CR *SI* reverses sign (indicating stimulus-selectivity) or remains similar (indicating outcome-selectivity). **Figure 3a** schematically plots the 5 major classes of *SI* changes that can be illustrated in a 2D before-after plot. Note that each neuron may show a mixture of stimulus- and outcome-selectivity, which are given by the projection components onto the two diagonals. We consider here only the major classes. To not only assess how neurons immediately react to the rule-switch but also how they may adapt during re-learning of the task, we assigned each neuron twice to these classes of *SI* changes (comparing *LE*→*RN* and *LE*→→*RE*, respectively; **Fig. 3a**). Among 107 chronically imaged lateral OFC neurons (n = 3 mice), we found a high fraction of outcome-selective neurons, which tracked outcome-value by responding strongly in the new-hit trials immediately following rule-switch in the *RN* period (**Fig. 3b-c**). Some OFC neurons also lost selectivity while others gained selectivity. This functional distribution was similar when comparing *LE* and *RE* periods (**Fig. 3d; Supplementary Fig. 9**). In S1, on the contrary, stimulus-selective neurons were more abundant than outcome-selective neurons in the *RN* period (18% of active neurons among 218 chronically imaged neurons; n = 4 mice; **Fig. 3e-f**). During reversal learning, however, the selectivity distribution of S1 neurons changed markedly. A large subpopulation had switched their selectivity in the *RE* period, compared to the *RN* period, functionally remapping to the new stimulus-outcome contingency (**Fig. 3g; Supplementary Fig. 9**). Moreover, a subpopulation of previously inactive or non-selective neurons acquired high outcome-selectivity for the newly rewarded go-texture. Identified S1→OFC projection neurons showed similar selectivity changes (**Supplementary Fig. 10**). A similar analysis of texture-touch-evoked responses in the stimulus-presentation window likewise revealed an overall remapping towards the new go-texture from the *RN* through the *RE* period (**Supplementary Fig. 11**). The link of functional subclasses to behavioural variables, especially reward-modulation of neurons with outcome-selectivity in *RE*, was further confirmed by GLM analysis^17^ (**Supplementary Fig. 12; Methods**). These results suggest that lateral OFC neurons exhibit a value-guided response immediately following a rule-switch. In contrast, a subpopulation of S1 neurons initially retains the learned stimulus-value association and is functionally remapped only upon re-learning.

**Figure 3.**
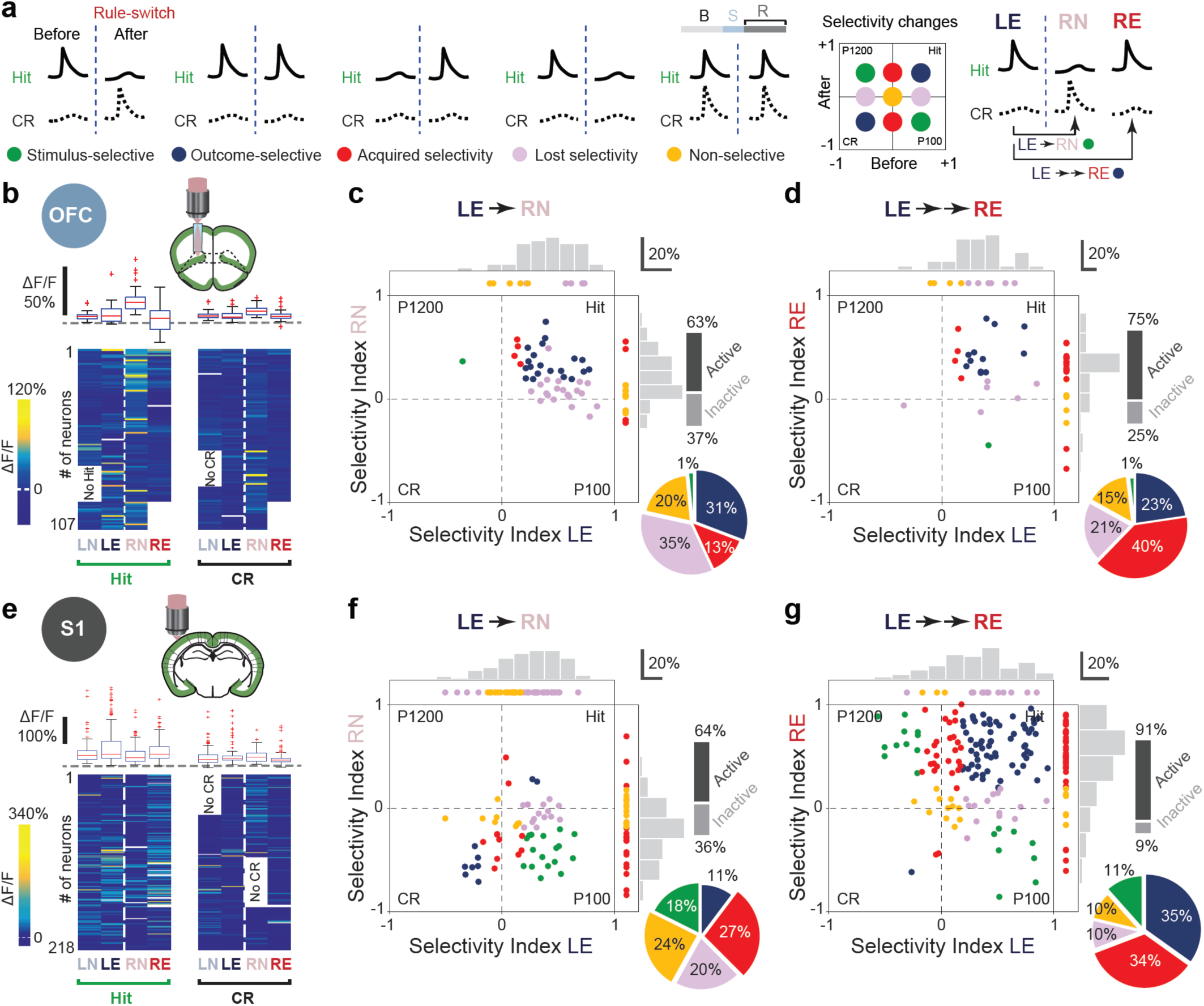
Distinct task-related functional dynamics in neuronal populations of lateral OFC and S1 after rule-switch. **a**, Schematic illustration of 5 major classes of hit/CR selectivity changes after rule-switch and their distribution in a 2D-scatter plot of selectivity before and after rule-switch. To the right, first- and second-class assignment for the *LE*→*RN and LE*→→*RE* comparisons. Selectivity was assessed using an ROC analysis. **b**, Mean ΔF/F amplitude in the reward-outcome window for lateral OFC neurons for hit (left) and CR (right) trials, averaged across each of the four salient phases. Bottom: Heat maps for 107 individual neurons, imaged longitudinally in three mice (n = 20 sessions). Top: Average values pooled across all neurons as box plots (red line: median, box edges: 25^th^ and 75^th^ percentiles, whiskers: minimum and maximum excluding outliers, red crosses: outliers outside 95% interval, dashed grey line: zero line). **c**, 2D-scatter plot and marginal distributions (histograms) comparing hit/CR selectivity of these lateral OFC neurons in *LE* and *RN* periods (selectivity index computed in reward-outcome window). Neurons that are only active in *LE* are displayed in the box above the 2D plot, neurons that are active in *RN* but not in *LE* are shown on the right. Active neurons with non-significant selectivity (*p* > 0.05, permutation test) are marked yellow. Note the high-fraction of outcome-selective OFC neurons. Neurons that are inactive in both phases are not included in the plot (percentage of active neurons shown on the right). **d**, Same plot as c but for *LE* versus *RE* period during which a fraction of lateral OFC outcome-selective neurons maintains the outcome-selective preference, while another subset of previously inactive neurons acquires new selectivity for the new-hit (n = 51 active neurons out of n = 68 chronically recorded in *LE* and *RE* in n = 3 mice, n = 16 sessions). **e**, Same plot as in b but for S1 neurons (n = 218 longitudinally recorded neurons in n = 4 mice, n = 28 sessions). **f**, Same plot as in c but for S1 neurons *LE* versus *RN* period, during which neurons retained their preference for the previous contingency (n = 90 active neurons over n = 142 chronically recorded in *LE* and *RN* in n = 4 mice, n = 20 sessions). **g**, Same plot as in f but for *LE* versus *RE* period, during which a subset of neurons updated their outcome-selective preference, and another subset of previously inactive neurons acquires new selectivity for the newly rewarded hit trials (n = 198 active neurons out of n = 218 chronically recorded neurons in *LE* and *RE* in n = 3 mice, n = 28 sessions).

Is the delayed remapping in S1 causally dependent on the activity in the lateral OFC? To investigate whether direct long-range OFC→S1 anatomical projections exist in mice, we injected retrograde AAV-retro/2-tdTomato into L2/3 of S1. Analysis of cleared brains (n = 2) using whole-brain light-sheet microscopy^18^ revealed dense S1-projecting neurons primarily in L2/3 and L5 of the lateral OFC (**Fig. 4a**). Chemogenetic silencing of lateral OFC neurons after the rule-switch (*RN* through *RE*) impaired functional remapping of S1 neurons (**Fig. 4b**; **Supplementary Fig. 9**; n = 4 mice). The effect is best seen in the marginal distributions for the three salient learning periods. Unlike control S1 neurons, a significant fraction of S1 neurons in OFC-silenced mice preserved their selectivity for the previous go-texture and failed to reconfigure responses towards the new go-texture in *RE*, with no significant difference observed in *LE* (cumulative distributions, two-sample Kolmogorov-Smirnov test) (**Fig. 4c**). Lateral OFC silencing also affected *RN*→*RE* remapping of texture touch-evoked Ca^2+^ responses in the stimulus-presentation window (**Supplementary Fig. 11)**. We additionally tracked neuronal fate by comparing the assigned classes for *LE*→*RN* and *LE*→→*RE* transitions. Whereas normally a fraction of non-selective and lost-selectivity S1 neurons (*LE*→*RN*) gained selectivity for the new go-texture (*LE*→→*RE*), we did not observe such recruitment for S1 neurons in lateral OFC-silenced mice (**Supplementary Fig. 9; Methods**). This analysis further confirms that S1 remapping crucially depends on top-down OFC input.

**Figure 4.**
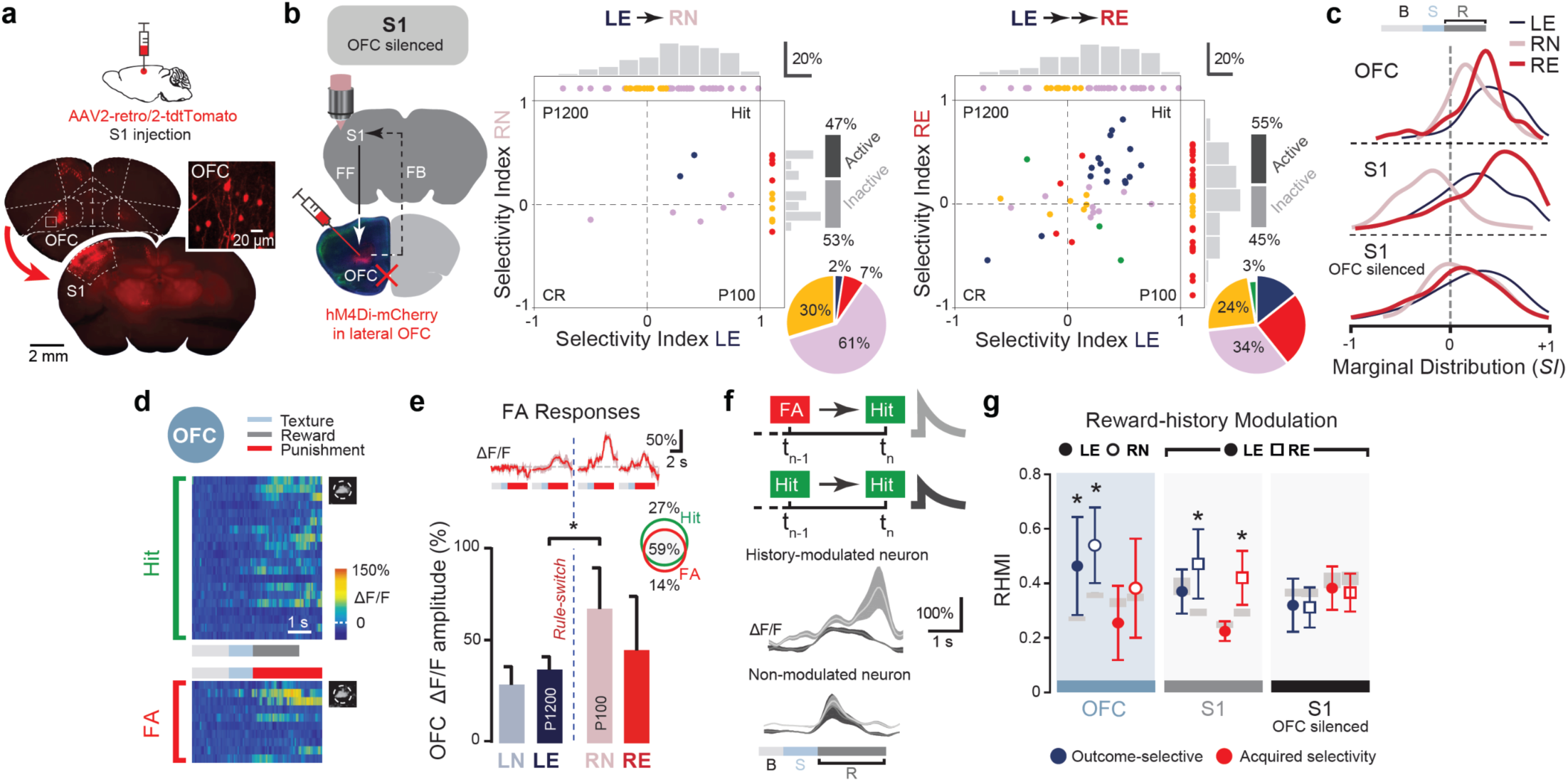
Long-range top-down lateral OFC input reconfigures functional responses of S1 neurons. **a**, Retrograde AAV-retro/2-tdTomato injection, CLARITY and whole-brain imaging revealed long-range lateral OFC→S1 feedback projections (n = 2 mice). *Inset*, layer L2/3 neurons in lateral OFC (scale bar = 20 μm). **b**, Left: L2/3 S1 neurons were chronically imaged while lateral OFC was chemogenetically silenced after the rule-switch. Middle and right: 2D-scatter plots of *SI* values computed in the reward-outcome window for *LE*→*RN* and *LE*→→*RE* together with marginal distributions as histograms (n = 164 chronically recorded neurons in *LE* and *RN*, n = 24 sessions, one session discarded due to motion artefact; n = 210 neurons longitudinally recorded in *LE* and *RE*, n = 25 sessions from n = 3 mice). **c**, Comparison of *SI* marginal distributions for the three salient periods *LE, RN*, and *RE* for OFC neurons (Fig. 3 c,d), S1 neurons (Fig. 3 f,g), and S1 neurons in lateral OFC-silenced mice (this figure, panel b). **d**, Heat-map of single-trial ΔF/F responses of an example lateral OFC neuron in the *RN* period sorted by hit and FA trials. Solid bars indicate different periods for texture-presentation (light blue), reward (grey), and punishment (white-noise, red). **e**, Average Ca^2+^ transients (top) and mean ΔF/F amplitudes (bottom) of FA trials for lateral OFC neurons during the four salient periods of learning and reversal learning (n = 63 active neurons out of n = 228 total neurons; n = 3 mice). *Inset*, Percentage of active neurons for hit and FA trials with overlap indicated. **f**, Averaged hit responses of two example outcome-selective neurons in S1 exhibiting trial-history dependent modulation with previous trial being rewarded (hit→hit; light grey trace) or punished (FA→hit; dark grey trace). **h**, Reward-history modulation index (RHMI) for outcome-selective neurons (blue) and neurons with acquired selectivity (red) neurons in the lateral OFC, S1, and S1 in OFC-silenced mice before (*LE*) and after (*RN, RE*) rule-switch (p < 0.05; bootstrap-permutation test; S.E.M. of RHMI with permutated indices as grey boxes).

Finally, we examined the mechanism of how lateral OFC governs S1 remapping by analysing the influence of error-modulation and history-dependence. Most lateral OFC neurons that responded strongly to new-hit trials also displayed enhanced response in FA trials in early phases of reversal (*RN*) indicating the error-dependent nature of OFC computation (**Fig. 4d-e**). Since OFC neurons integrate recent reward-history^22^, we also studied intrinsic features of value-sensitive neurons in both lateral OFC and S1. We computed a ‘reward-history modulation index’ (RHMI) for each neuronal subclass by comparing responses of each of the ‘hit’ trials that were immediately preceded by a ‘hit’ or an ‘FA’ trial (**Fig. 4f**; **Methods**). While outcome-selective neurons in the lateral OFC showed a significant reward-history-dependent response modulation before (*LE*) and after (*RN*) the rule-switch, S1 neurons with both outcome-selectivity and acquired-selectivity, but no other subclasses, were modulated by reward-history in *RE*. Interestingly, history-dependent modulation was absent in S1 neurons in OFC-silenced mice indicating that higher-order top-down input from the lateral OFC is critical for the intrinsic functional reorganisation of both outcome-selective and gain-selectivity neurons in S1 (**Fig. 4g**; **Supplementary Fig. 13**). These findings corroborate the notion that encoding of outcome-value in the lateral OFC, as well as deviations from expected outcome-value, is critical for functional remapping of a selective subpopulation of S1 neurons for flexible decision-making.

Adaptive behaviour is shaped by recent and accumulated sensory evidences, as well as predicted outcome-value of future choices. Predictive processing is critical for perception^19^ and the OFC provides a ‘cognitive map’ functioning as an internal reference framework to compare and optimise a desired behavioural outcome such as reward^20,21^. Reward value is computed in distributed parts of the brain^22^ including the VTA^23^, frontal areas^7^, and primary and associational^17^ sensory areas of the neocortex. Our experiments revealed a crucial role of mouse lateral OFC neurons in encoding outcome-value prediction error as a teaching signal for driving learning of an altered stimulus-reward contingency, partly resembling classical dopamine responses^23,35^. Tracking both positive and negative outcome-values, OFC neurons may represent ongoing neural estimates of position on a value map^24^. Pharmacogenetic silencing revealed that lateral OFC has a critical causal role in implementing flexibility as previously shown in rodents^25^, as compared to non-human primates^26^. This implementation could be achieved via possible projection-specific interactions^27^ with integrative cortical areas like the retrosplenial cortex^28^, and/or subcortical structures including the basolateral amygdala^27,29^ and the mediodorsal thalamus^30^. Our findings also crucially imply that S1 neurons do not simply function as sensory feature detectors^31^. Whereas reward-sensitive neurons were to be expected in the higher-order cortex such as OFC, perhaps more importantly, we discovered a subpopulation of neurons in the primary sensory cortex that are differentially modulated by reward-history. The cellular and circuit mechanisms for a remarkable plasticity in this small subset of neurons, possibly involving neuromodulators such as serotonin^32^ or long-range layer-specific excitatory and local inhibitory interactions^33^, remain to be determined. The existence of a reward-valence signal in the primary sensory cortex and their modulation by higher-order inputs has important implications for implementing reinforcement learning algorithms for brain machine interfaces^34^. Taken together, this study revealed crucial cellular-level local interactive motifs and long-range circuit interactions behind adaptive plasticity and flexible decision-making.

## Acknowledgements

This work is supported by a H2020 Marie Sklodowska-Curie fellowship (CIRCDYN, Grant number: 709288) and a NARSAD Young Investigator award (Grant number: 24941) from the Brain & Behavior Research Foundation (to A.B.), grants from the Swiss National Science Foundation (Grant number: 310030B_170269), and the European Research Council (ERC Advanced Grant BRAINCOMPATH, Grant number: 670757) grant (to F.H.). The authors thank Prof. Benjamin Grewe (ETH-Zürich) for showing us optical preparation involving GRIN lens, and Prof. Martin Schwab for the use of equipment for open-field and ladder-rung test. The authors thank Stefano Carta, Lazar Shumanovski, Dubravka Göckeritz, Dr. Ladan Egolf, Chiara Rickenbach for their various assistance. The authors thank Dr. Walter Senn, Dr. Federica Lucantonio, Dr. Michael Goard, Dr. Benjamin Scholl for helpful discussions on the manuscript. The authors declare no conflict of interest.

## Author contributions

A.B. conceived the project. A.B. and F.H. designed the study. A.B and J.T. carried out all experiments except multi-unit electrophysiology experiments (performed and analysed by C.L.). A.B and G.P. analysed data. F.F.V developed light-sheet microscope and imaged cleared brains together with A.B. A.B. and F.H. wrote the manuscript with comments from all authors.

