## Supplementary Material for "Value-guided remapping of sensory circuits by lateral orbitofrontal cortex in reversal learning"

#### Methods

**Animals.** All experimental procedures were carried out in accordance with the guidelines of the Federal Veterinary Office of Switzerland and were approved by the Cantonal Veterinary Office in Zurich under license numbers 285/2014 and 234/2018. A total of 30 adult male mice (6-8-week old) were used in this study. For behavioural experiments, we used wild-type (WT) C57BL6/J mice (n = 16 mice). For imaging neurons in lateral OFC and S1, we used Rasgrf2-2A-dCre: CamK2a-tTA: TITL-GCaMP6f triple transgenic mice, expressing GCaMP6f in excitatory neocortical layer 2/3 neurons (n = 14 mice). For causal pharmacogenetic manipulations, both WT and L2/3-GCaMP6f animals were used (n = 3 WT mice and n = 3 GCaMP6f mice). To generate triple transgenic animals amenable to two-photon imaging, double transgenic mice carrying CamK2a-tTA (JAX# 016198[36]) and TITL-GCaMP6f (JAX# 024103[37]) were crossed with a Rasgrf2-2A-dCre line (JAX# 022864[38]). The de-stabilised Cre-recombinase expressed under the control of the Rasgrf2-2A promoter was stabilised by trimethoprim (TMP, Sigma T7883) to render it functional. TMP was reconstituted in Dimethyl Sulfoxide (DMSO, Sigma 34869, 100 mg/ml), freshly prepared before each induction, and administered two weeks before surgery. During induction, mice were given a single intraperitoneal injection (150 mg TMP/g body weight diluted in 0.9% saline solution) using a 29g needle. To specifically label and image from S1→OFC projection neurons, we injected AAV2.9.hSyn.FLEX.GCaMP6f virus into S1 of WT mice. Mice were grouped with their WT siblings and housed at 24°C and variable humidity in 12-hour reverse dark-light cycle (7:00 a.m. to 7:00 p.m.). At the end of an experiment, the animals were deeply anesthetised and transcardially perfused or euthanised with an overdose of pentobarbital (150 mg/kg body weight, i.p.). All efforts were made to minimise suffering. All mice belonged to the C57BL6/J strain.

**Reversal learning task.** Mice were extensively handled during pre-training sessions to familiarize them with the experimenter and experimental setup. Once they had acclimatised to handling, mice were placed on water-restriction and trained on a go/no-go tactile-discrimination task. Mice remained on water-restriction for the remainder of the experiment. The behaviour set-up has been described previously[10]. During the start of each trial, an auditory cue (2 beeps at 2 kHz, 100 ms duration with 50 ms interval), indicated the approach of one of two possible textures (sandpapers of grit size P100, rough texture; P1200, smooth texture). The texture was positioned to reach the mouse's whiskers and 'go' or 'no-go' textures were presented pseudorandomly with no more than three consecutive repetitions. The texture stayed in touch with the whiskers for one second (‘sensation’), after which it moved out of reach. An additional auditory tone (response cue; 4 beeps at 4 kHz, 50-ms duration with a 25-ms interval) signalled the start of a 2-second ‘response window’ during which the mouse had to lick or withhold from licking the water sprout to indicate its choice (‘outcome or response’, 2 seconds). A sucrose-water reward was delivered only for licks in response to the ‘go’ texture and after the response cue (‘hit’). Incorrect licks in response to the non-target ‘no-go’ texture (‘false alarms’, FA) was punished with a brief period of mild auditory white noise. Reward and punishment were omitted when mice withheld licking for the no-go (‘correct-rejections’, CR) or go (‘miss’) textures. The licking detector remained in a fixed and reachable position throughout the entire trial.

Mice proficiently performed the sensory-discrimination task from learning naïve (*LN*) through expert phase (*LE*). Once mice had achieved stable performance of the tactile-discrimination task (reaching a $d' = 1.5$ for 3-4 sessions), the stimulus-response mapping was switched (‘rule-switch’). Upon rule-switch, performance initially dropped to chance level or below. However, after 4-5 days, all mice (n = 11 out of 11 mice) learned the new texture-response mapping and increase performance from reversal naïve (*RN*) through expert phase (*RE*) as quantified by the increase in the discriminability index (*d’*) (training period 4-5 days, 200-300 trials/session/day).

*Animal training and performance measurement.* We quantified mice task performance using the discriminability index d-prime (*d’*) rather than percent correct to account for motivation and criterion[39]. We set the learning threshold to $d' = 1.5$. $d'$ was calculated for each session as $= Z(\text{hit}/(\text{hit+miss})) - Z(\text{FA}/(\text{FA+CR}))$ with $Z(p)$, $p \in [0,1]$, being the inverse of the cumulative Gaussian distribution (FA, number of false alarm trials; CR, number of correct rejection trials). We selected in both training periods pre- and post-reversal two relevant phases corresponding to the salient phases - learning and reversal naïve (*LN* and *RN*, respectively), in which the mice were performing lower or close to chance level ($d' = 0$, $p < 0.05$ for $d' > 0$, n = 1-3 sessions), and learning and reversal expert (*LE* and *RE*, respectively, n = 1-3 sessions), in which the mice were stably performing above a criterion set as $d' = 1.5$. Expert sessions were always selected from the last sessions available immediately before rule-switch (*LE*) or task completion (*RE*), and this resulted in high performance level ($d' > 2$). For imaging data, only days among these respective sessions were used.

**Whisking and licking measurement.** During task performance, whisker kinematics and fine body movement were simultaneously monitored using high-speed cameras. We identified behavioural correlates of task learning by quantifying licking rate and whisking amplitude obtained from lick-sensor measurements and high-speed videography, respectively. The whiskers were illuminated with 940 nm infrared LED light and movies were acquired during the behaviour at 500 Hz (500 × 500 pixels) using a high-speed CMOS camera (A504k; Basler). Average whisker angle across all imaged whiskers was measured using automated whisker tracking software. The whisking amplitude (envelope) was calculated as the difference in maximum and minimum whisker angle along a sliding window equal to the imaging frame duration (142 ms). Principal whisker velocity was calculated by applying a band-pass filter to the whisking angle time vector and then computing its first derivative. For all trials recorded (n = 3 mice), the first and last possible time point for whisker-to-texture contact was quantified manually through visual inspection.

Licking was detected by using a piezo-electric sensor attached to the lick spout and lick rates were calculated by thresholding this signal and counting the number of events per unit of time. Multiple consecutive threshold crossings which occur in rapid succession can result in a lick rate that exceeds the physical capability of a mouse. We therefore made the reasonable assumption of a peak lick rate of 10 Hz based on manual checks on videography. A low pass filter was applied to the lick rate time series, which effectively combined multiple events occurring within a 100 ms period into one event. Expert mice showed a decrease of early licks. While early licks are not exhibited immediately upon rule-switch when the behavioural performance is low, lick rates are slightly lower compared to expert sessions.

**Open-field test.** General locomotor activity was measured in an open-field (a rectangular arena of 40 x 30 x 20 cm)[40] made from grey Plexiglas that was illuminated from a centred diffuse light source. A single animal was exposed to the environment for 5 minutes while being recorded by a video camera placed above the open field and operated by LabVIEW (National Instruments). Mouse velocity (cm/s) and distance covered (cm) were analysed using the EthoVision software.

**Horizontal ladder-rung test.** A 1-m long horizontal ladder, consisting of two platforms connected by an irregular pattern of 70 rungs was used. The distance between rungs varied between 1-3 cm. Mice were given time to practice with three trials before being tested. Three trial sessions per animal were recorded using a high-speed camera (Nikon AF Nikkor) at 100 frames per second. Each forepaw placement was analysed and the quality of the placement was scored using the following scoring system[41]. A perfect paw placement on the rung was scored as 1; partial digit placement, correction and replacement were scored as 0.5, slip or total miss were scored as 0. The success rate was calculated for each animal group as

$$\textit{Success rate} = (\textit{Total score}/\textit{Number of steps}) \times 100 \quad (1)$$

**Virus injection.** Mice were briefly anaesthetised with isoflurane (2%) in oxygen in an anaesthesia chamber and subsequently transferred to a stereotactic frame (Kopf Instruments). Body temperature was maintained at ~37°C using a heating blanket with a rectal thermal probe. The eyes of the mouse were covered by Vitamin A cream (Bausch & Lomb) during the surgery. The cranium was secured with ear bars and anaesthesia was maintained during the surgery with 0.8-1.2% isoflurane. After disinfection with Betadine, the skin was opened using a scalpel and an L-shaped incision was made in the skin, and the cranial surface was cleaned using absorbent swabs (Sugi; Kettenbach GmbH). We identified lateral OFC based on stereotactic coordinates from previous studies (2.6 mm anterior and 1.2 mm lateral from bregma)[13]. For S1, injection coordinates were 3.5 mm lateral and 1.5 mm posterior from bregma. The skull was thinned along a 1-mm line at the rostral edge of S1 using a Dremel drill with occasional cooling with saline. After drilling through the cranium, the dura was punctured using a glass micropipette filled with the virus suspended in mineral oil. Several injections (3-4) were made at neighbouring sites, at a depth of 200-250 µm. A volume of 100-150 nl of virus was injected at 50 nl/min rate at each site. After each injection, the pipette was held in place for 5-8 minutes before retraction to prevent leakage. Skin was sutured using a synthetic, monofilament, non-absorbable suture (Prolene 7.0, Ethicon).

**Cranial window and GRIN lens implantation.** To study neural dynamics in the lateral OFC, a chronically implanted metallic cannula was implanted on top of lateral OFC with a glass coverslip at its base. Cannula implantation and cranial window preparation was performed under isoflurane anaesthesia following details as described above. A circular piece of cranial bone (diameter ~ 1.5 mm) was drilled on top of OFC using a Dremel drill. A modified biopsy punch (diameter 1.0 mm; Miltex) was inserted 1.5 mm deep into the cortical tissue for two minutes. The cortical tissue (primary and secondary motor areas) was gently aspirated with a cut using a 27-gauge needle connected to a water jet pump, while constantly being rinsed with Ringer solution. We removed the overlying cortex using aspiration until layer 5 (depth

1.5-1.7 mm) and implanted a stainless-steel cannula (internal diameter 1.0 mm, 1.5 mm height) was with its base covered by a cover glass (0.17 mm thickness) 1.6-1.8 mm below the pial surface. The cannula was secured in place by UV curable dental acrylic cement (Ivoclar Vivadent). We waited two-three weeks after surgery before commencing training. Before each imaging session, a rod-like gradient-index (GRIN) lens (NEM-100-48-00-50-NC, customised needle endomicroscope for two-photon microscopy, ~ 0.4 pitch, corrected for wavelength $\lambda$ = 920 nm, diameter = 1.0 mm, length ~ 4.3 mm; GRINTECH GmbH, Jena) was inserted through the cannula and neurons were imaged 100-300 um below. Before each imaging session, the cannula was cleaned with distilled water.

To allow long-term *in vivo* calcium imaging in S1, a cranial window was implanted over S1 as described previously[10,42]. A metallic head-post for head fixation was glued to the skull, contralateral to the cranial window, using dental acrylic. One week after chronic window implantation, mice were handled daily for one week while they became acclimatised to a minimum of 15 mins of head-fixation.

**Brain clearing and light-sheet microscopy.** To verify task-relevant projections and connectivity between S1 and lateral OFC, we injected retrograde AAV-retro/2-shortCAG-tdTomato virus *in vivo*. Two to three weeks after virus injection, animals were perfused, and the brains entered a clearing protocol using CLARITY[43]. After perfusion, the brains were post-fixed for 48 hours in a hydrogel solution (1% paraformaldehyde, 4% acrylamide, 0.05% bis-acrylamide, 0.25% VA044)[43,44] before the hydrogel polymerization was induced at 37°C. Following the polymerization, the brains were immersed in 40 ml of 8% SDS and kept shaking at room temperature (RT) until the tissue was cleared sufficiently (20-40 days depending on the age of the animals). Finally, after 2-4 washes in PBS, the brains were put into a refractive index matching solution (RIMS)[43] for the last clearing step. They were left to equilibrate in 5 ml of RIMS for at least 4 days at RT before being imaged.

Cleared brains were imaged using a mesoSPIM light-sheet microscope (www.mesospim.org)[18]. Whole-brain imaging revealed that lateral OFC receives direct monosynaptic bottom-up, feed-forward projections from both superficial (L2/3) and mostly deep (L5 and L6) layers of S1. Conversely, a similar injection in mouse S1 (2.55 mm posterior and 3.5 mm lateral from bregma)[10] revealed superficial cortical L2/3 neurons in mouse S1 receiving direct top-down feedback projections from lateral OFC.

**CNO application.** Inhibitory DREADDs (CaMKIIα-hM4D($G_i$)-mCherry) were used in the chemogenetic silencing experiments and neuronal populations of interest were virally transfected with AAV-hM4Di injected unilaterally on the superficial layers (L2/3) of contralateral OFC and bilaterally to superficial (L2/3) and deeper (L5) layers of S1. Intraperitoneal (i.p.) injection of clozapine-N-oxide (CNO, 1-5 mg/kg), the ligand that activates hM4Di, silenced the activity of L2/3 and L5 neurons. Clozapine (1-5 mg/kg) was used as control[45].

***In vivo* electrophysiological recordings.** We characterised pharmacogenetic silencing of lateral OFC neurons by performing acute, *in vivo* electrophysiology in a subset of hM4Di-injected animals after completion of the reversal learning protocol. To perform acute recordings, animals were anesthetised with isoflurane (2% for induction and 0.8% during recording), and their body temperature was maintained stably using a heating pad. A small craniotomy (1-mm diameter) was performed to provide access to the left OFC and the brain was covered with silicon oil. A silver wire was placed in contact with the CSF through a small trepanation (0.5 mm) over the cerebellum to serve as reference electrode. A silicon probe (Atlas Neurotechnologies, 16 linear sites, 100 µm spacing) was implanted through the craniotomy into the left cortical hemisphere and we recorded multi-unit activity from the injection site in the left OFC and surrounding cortex. We waited 30 minutes to allow the recording to stabilise after implantation of the electrode array. After stabilisation, the broadband voltage was amplified and digitally sampled at a rate of 30 kHz using a commercial extracellular recording system (RHD2000, Intan Technologies). The raw voltage traces were filtered offline to separate the multi-unit activity (MUA; bandpass filter 0.46-6 kHz) using a fourth-order Butterworth filter. Subsequently, the high-pass data were thresholded at 6.5 times the standard deviation across the recording session and the numbers of spikes in windows of interest were counted. After a baseline recording of 30 mins, CNO (1-5 mg/kg) was injected (i.p.). During the baseline period (30 minutes), the average firing rate remained stable, while upon CNO injection the average firing rate in the lateral OFC steadily decreased over time. Recording electrodes in the lateral OFC showed a stable and significant decrease in spiking activity 30 minutes after CNO administration, while control electrodes from areas uninfected by the virus did not show any modulation. To combine data across mice, the activity at sites with clear MUA was expressed in percent of the baseline value, i.e. the average spike rate during the 30-minute pre-injection baseline (100%). All multi-units were then combined from the injected or control region and a t-test was performed between the baseline period (-30-0 minutes pre-injection) and the post injection period (30-60 minutes post injection).

**Intrinsic signal optical imaging.** The S1 barrel cortex was identified using intrinsic signal optical imaging under approximately 0.8-1 % isoflurane anaesthesia. The cortical surface was illuminated with a 630-nm LED, multiple whiskers were stimulated (2 to 4 rostro-caudal deflections at 10 Hz), and reflectance images were collected through an objective with a CCD camera (Toshiba TELI CS3960DCL; 12-bit; 3-pixel binning, 4273 347 binned pixels, 8.6-mm pixel size, 10-Hz frame rate)[46].

Intrinsic signal changes were computed as fractional changes in reflectance relative to the pre-stimulus average (50 frames; expressed as DR/R). The centres of the barrel columns corresponding to stimulated whiskers were located by averaging intrinsic signals (15 trials), median-filtering (5-pixel radius) and thresholding to find signal minima. Reference surface vasculature images were obtained using 546-nm LED and matched to images acquired during two-photon imaging.

**Two-photon imaging.** We used a custom-built two-photon microscope controlled by HelioScan[47], equipped with a Ti:Sapphire laser system (approximately 100-femtosecond (fs) laser pulses; Mai Tai HP; Newport Spectra Physics), a water-immersion 16X Olympus objective (340LUMPlanFI/IR, 0.8 numerical aperture, NA) for S1 imaging and a 20X Leica objective (Leica Plan Apo 0.6 NA) for GRIN lens based OFC imaging, galvanometric scan mirrors (model 6210; Cambridge Technology), and a Pockels Cell (Conoptics) for laser intensity modulation.

Based on intrinsic imaging, along with the blood vessel pattern, we targeted specific areas of interest for two-photon imaging of L2/3 neurons in each mouse. We excited GCaMP6f at 940 nm and detected green fluorescence with a photomultiplier tube (Hamamatsu). Images (128x64 pixels) were acquired at 12-Hz frame rate and 10-50 cells per field of view were imaged simultaneously. Single trials of 6-8 s duration were recorded, with 1-s breaks between trials to allow the data to be written to hard-disk during inter-trial periods.

**Calcium imaging analysis.** Calcium imaging data was first motion corrected using an online piecewise rigid 2d (planar) method (NoRMCorre: Non-Rigid Motion Correction) in MATLAB (Mathworks). Regions of interest (ROI) corresponding to individual neurons were found from both the mean image and the standard deviation image generated from a single-trial time series using ImageJ (US National Institutes of Health). ROI masks were manually selected using an online method (OCIA) in

MATLAB and raw fluorescence time courses (F(t)) were then extracted as the (non-weighted) mean pixel value for each ROI. Another fluorescence time course was extracted from a neuropil defined by an ROI selecting a portion of non-somatic tissue in the imaging frame. The neuropil calcium signal never resulted in activity peaks significantly high to be classified as an active neuron (check Criteria for active neurons). The background was subtracted on each channel (bottom first percentile fluorescence signal across entire time series). A running estimate of fractional change in fluorescence time courses was calculated by subtracting the baseline fluorescence $F_0(t)$ from F(t), then dividing by $F_0(t)$

$$\Delta F/F(t) = (F(t) - F_0(t))/F_0(t) \qquad (2)$$

$F_0(t)$ was estimated as the mean fluorescence value of the first 1.5 s prior to tactile stimulus onset. For cells that were not silent in the pre-stimulus period, $F_0(t)$ was instead taken as $8^{th}$ percentile of a trailing 1.5-s sliding window.

**Alignment of cell-masks across days.** All analyses for the alignment of cell-masks across days were manually performed with the aid of custom MATLAB GUIs in the OCIA software. To align masks across any pair of daily sessions, we first chose one set for the first day and then imported it onto the single-trial image series of the subsequent days. When displacement occurred, the masks were manually moved to the corresponding neurons. This was done for all pairwise combinations of days. We then manually observed by eye each ROI mask confronting it to both the mean and the standard deviation image of the time series on ImageJ, to confirm the presence of each cell across days. If the z-plane did not match and a cell was not found, it was excluded from further longitudinal analysis.

**Criteria for active neurons.** To determine if a neuron was active during a time-period of interest (stimulus-related and reward-outcome related responses), we independently tested its evoked response using conservative criteria. For each neuron, we calculated its mean response and its peak value ($\Delta F/F$) during the 0.9 s period after a texture was presented (i.e. for stimulus presentation-window) or during the1.6-s period after the texture was removed (i.e. for reward-outcome window). A neuron was considered active if all the following criteria were met:

- its response was significantly ($p < 0.01$, *t*-test) different from the average pre-stimulus baseline response (1.5 s before texture is presented).

- its mean response (for stimulus-presentation or reward-outcome window) was more than 3*noise from the baseline. This baseline was calculated by averaging a 35-point sliding-window across the trial response and taking the $5^{th}$ percentile of the mean response distribution. The noise level taken as the $1^{st}$ percentile of the distribution of the standard deviation calculated across the same sliding window.
- its peak response (ΔF/F) (for stimulus or reward-outcome window) was greater than 25%.
- In the 2D scatter plots of selectivity indices (see below) neurons were considered active if they were active in either of the considered learning periods (e.g. *LE* and *RN*). In other words, they were considered inactive only if they were inactive in both respective periods.

**Selectivity index.** We assessed the selectivity of single-neuron activity for specific trial-types using a receiver operating characteristic (ROC) analysis, which quantifies the ability of an ideal observer to discriminate between trial-types based on single-trial responses[16,10]. For this study, we assessed selectivity for hit vs. CR trials. We performed the ROC analysis on the segments of the ΔF/F transients in the trial period of interest, i.e., either in the 2 s long reward-outcome window or in the 1 s long stimulus window. Specifically, each trial was assigned a "discrimination variable" score (*DV*) equal to the dot product similarity of the ΔF/F segment to the mean ΔF/F segment for the same trial-type minus the dot-product similarity to the mean for the other trial-type (see also **Supplementary Fig. 8**). Thus, we computed for hit trials

$$\boldsymbol{DV_{Hit}} = \boldsymbol{H_i}(\boldsymbol{\overline{C}_{\forall j \neq i}} - \boldsymbol{\overline{C}}) \quad (3)$$

and for CR trials

$$\boldsymbol{DV_{CR,i}} = \boldsymbol{C_i}(\boldsymbol{\overline{H}} - \boldsymbol{\overline{C}_{\forall j \neq i}}) \quad (4)$$

where $H_i$ and $C_i$ are the single-trial ΔF/F segments for the *i*-th hit and CR trial, respectively, and $\boldsymbol{\overline{H}}$ and $\boldsymbol{\overline{C}}$ denote the mean ΔF/F segments for the respective trial type (excluding the individual trial under consideration). We classified trials as belonging to the go-texture or the no-go-texture if *DV* ($DV_{Hit}$ or $DV_{CR}$) was greater than a given criterion. To determine the fraction of trials an ideal observer could correctly classify, we constructed an ROC curve by varying this criterion value across the range of *DV*. At each criterion value, we plotted the probability that a hit trial exceeded the criterion value against the probability that a CR trial exceeded the criterion value. The area under this ROC curve (AUC) indicates the selectivity for trial type, with an AUC value of 0.5, meaning no selectivity. We defined the "selectivity index", *SI*, such that it spanned the range from -1 (CR-preferring neurons) to +1 (hit-preferring neurons) by calculating

$$\boldsymbol{SI} = \mathbf{2}\times(\boldsymbol{AUC} - \mathbf{0.5}) \qquad (5)$$

We tested whether neurons showed trial-type selectivity above chance using a permutation test creating 500 permutations with trial-type labels randomly shuffled. From these permutations, we created a distribution of indices that could have arisen by chance and considered a neuron's *SI* value as significant if it fell outside the centre 95% interval of this distribution ($p < 0.05$).

**Functional classification of neurons.** Neurons that met the activity criterion in at least one of the salient learning periods were classified in different groups according to their hit/CR *SI* value changes upon rule-switch. For each of these neurons we compared the *SI* value in the pre-reversal period (*LE*) to the *SI* value in the two post-reversal periods (*RN* and *RE*). This resulted in two classifications for each neuron (for *LE→RN* comparison and *LE→→RE* comparison) (**Fig. 3a**). When two *SI* values before and after reversal were found concordant, i.e. of the same sign and significant, a neuron's response was classified as 'outcome-selective' for the respective post-reversal phase and the specific trial time-window considered (stimulus or reward-outcome). Such neuron's response amplitude was significantly higher for hit compared to CR trials (or CR compared to hit trials) independent of stimulus-identity (in the 2D scatter plots these neurons are found in the upper right and lower left quadrants). When *SI* values before and after reversal were discordant, i.e., of opposite sign and significant, the neuron's response was classified as 'stimulus-selective' as it switched from hit- to CR-preferring (or CR- to hit-preferring), where the new CR was associated with the same stimulus as the previous hit. In the 2D scatter plot these neurons are found in the upper left and lower right quadrants. If an active neuron was discriminating above chance during the pre-reversal period *LE* and lost significant selectivity in the pre-reversal period considered (*RN* or *RE*), or if it simply became inactive, it was classified as a 'lost-selectivity' neuron. Likewise, if an inactive neuron or an active neuron without significant selectivity in the pre-reversal period became active and gained a significant selectivity for the new hit/CR trials, it was included in the 'acquired-selectivity' group. Finally, all the active neurons that did not show a significant *SI* value during either phase (based on permutation tests), were considered 'non-selective'. Each one of these neurons was assigned twice to a functional group, in earlier (*RN*) and later phases of reversal (*RE*). We tracked the class transition through the course of re-learning using a fate map. For each *LE→RN* group we showed the fraction of neurons falling into the new *LE→→RE* classes. Only active neurons during both phases are shown.

**Reward-history modulation index.** To quantify the effect of previous performance on neural responses, we analysed how response magnitude varied as a result of the outcome of the previous trial (punishment or reward)[17]. We compared the response magnitude of each neuron during a hit trial when the previous trial was rewarded hit ($R_{Hit-Hit}$) versus the response magnitude when the previous trial was punished ($R_{FA-Hit}$). To quantify modulation by previous trial history, we created a reward-history modulation index (RHMI) by normalizing the difference between these two history-dependent responses by the mean overall response of all the Hit trials:

$$RHMI = \frac{|R_{FA-Hit} - R_{Hit-Hit}|}{R_{Hit}} \quad (6)$$

Only cells that were active during a specific phase were included in the RHMI analysis for that respective phase. To check whether a neuron was modulated above chance, a bootstrap permutation test was performed (500 permutations).

**Generalized linear model.** To estimate the contribution of behavioural and task variables (cue, stimulus onset and offset separated by behavioural response, reward delivery, punishment, licking) to the activity of each neuron, we fit a Poisson generalized linear model (GLM) for each session (MATLAB glmnet package). We first down-sampled deconvolved neural data and all behavioural and task variables to 10 Hz and then smoothed neural activity using a Gaussian filter. Regression functions were created from behavioural and task variables by implementing vectors of Gaussian filters (all filters had a standard deviation of 1 s, overlapping and evenly distributed, 1 Gaussian/3 frames, 100 ms/frame, 144 filters). Each imaging session consisted of 100-120 trials of 6 seconds each (15 Hz) (training set 75% of each run, testing sets 25%; 10-fold cross validated with 11 evenly spaced chunks of trials). We used an elastic net regularization consisting of 99% L2 and 1% L1 methods for each individual neuron. Deviance explained was calculated by comparing the activity predicted by the model to the actual activity calculated using data not used during the fitting procedure. Finally, the contribution of each variable to the neural activity is derived by calculating again the deviance explained using just that variable and normalizing it to the total deviance explained. This is plotted separately for each group of neurons.

**Statistical analysis.** Statistical analyses are described in the main text and in figure legends. In general, we used non-parametric statistical analyses (Wilcoxon sign-rank test, rank-sum test) or permutation tests to avoid assumptions about the distributions of the data. All statistical analysis was performed using custom written routines in MATLAB. When assumptions could be made based on previous literature, *t*-test was used. Quantitative approaches were not used to determine if the data met the assumptions of the parametric tests.

Suppl. Fig. 1

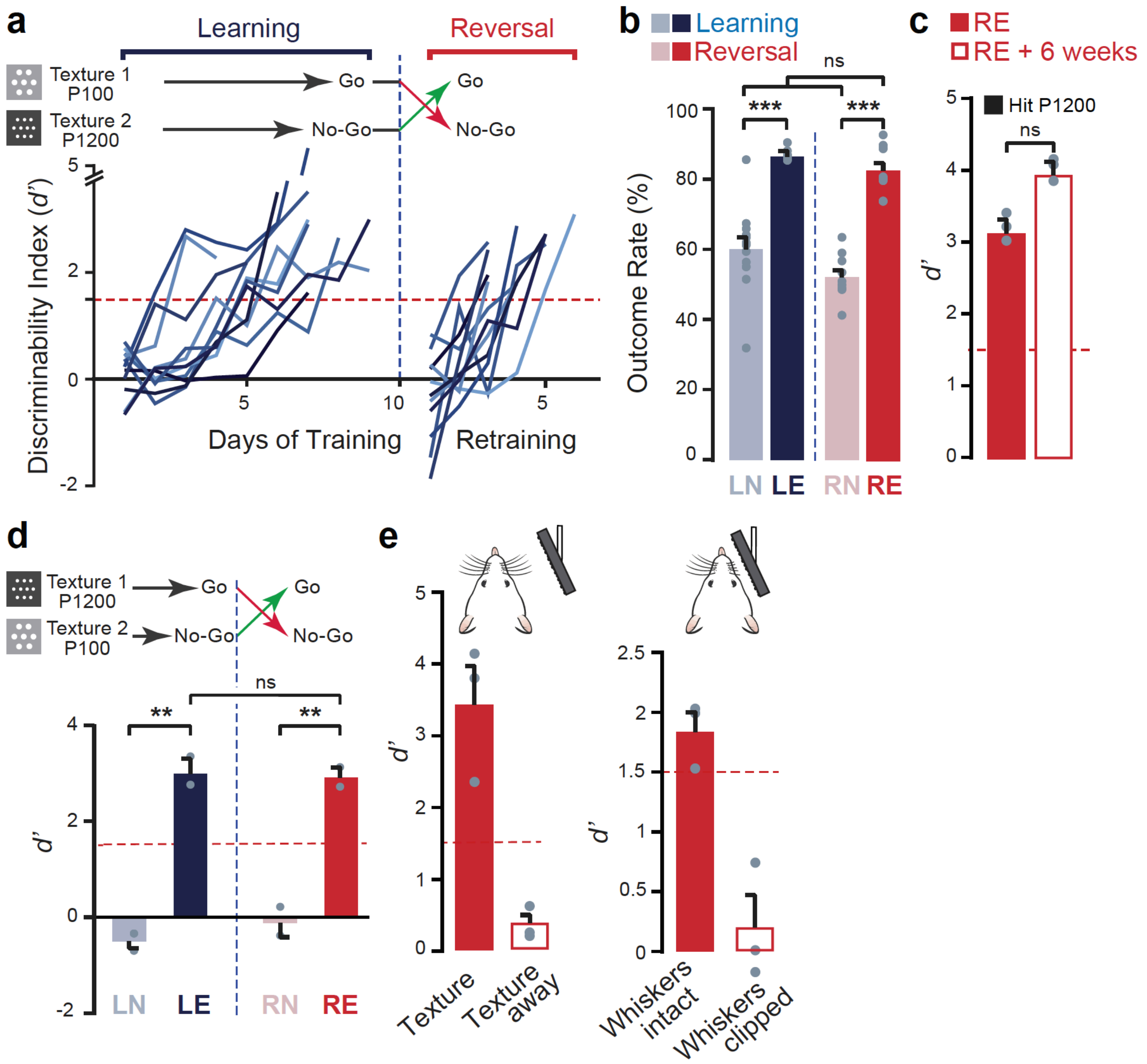


**Supplementary Figure 1 | S1-dependent tactile-discrimination-based reversal learning task. a,** Time-course of task-performance (discriminability index, *d'*) of individual mouse reveals dynamics of learning and reversal learning upon rule-switch (dashed blue-line). Each colour represents a single mouse of a total of 11 mice. **b,** Percentage of correct decision '(hit+CR)/all trials' as 'outcome rate' plotted during the four salient behavioural phases of learning (learning naïve, *LN*; learning expert, *LE*) and reversal (reversal naïve, *RN*; reversal expert, *RE*) (n = 11 mice). **c,** Reversal performance is stable and remains high when mice with reversed reward contingency (P1200 as go-texture, *RE*) were tested six weeks later (n = 2 mice). **d,** Reversal learning is independent of initial texture training (fine grit size sandpaper P1200 as initial go-texture; n = 2 mice). **e,** Texture-discrimination is dependent on sensory input. *Left:* Keeping textures out of reach in expert mice after reversal (*RE*) impaired their performances (n = 3 sessions in 2 mice). *Right:* Clipping whiskers in expert mice similarly resulted in impaired performance (low *d'*) indicating sensory input is essential for the correct execution of the task (n = 3 mice, longitudinally studied before and after whisker-clipping). All data presented as mean ± S.E.M., *p < 0.05, **p < 0.01 Wilcoxon rank-sum test.

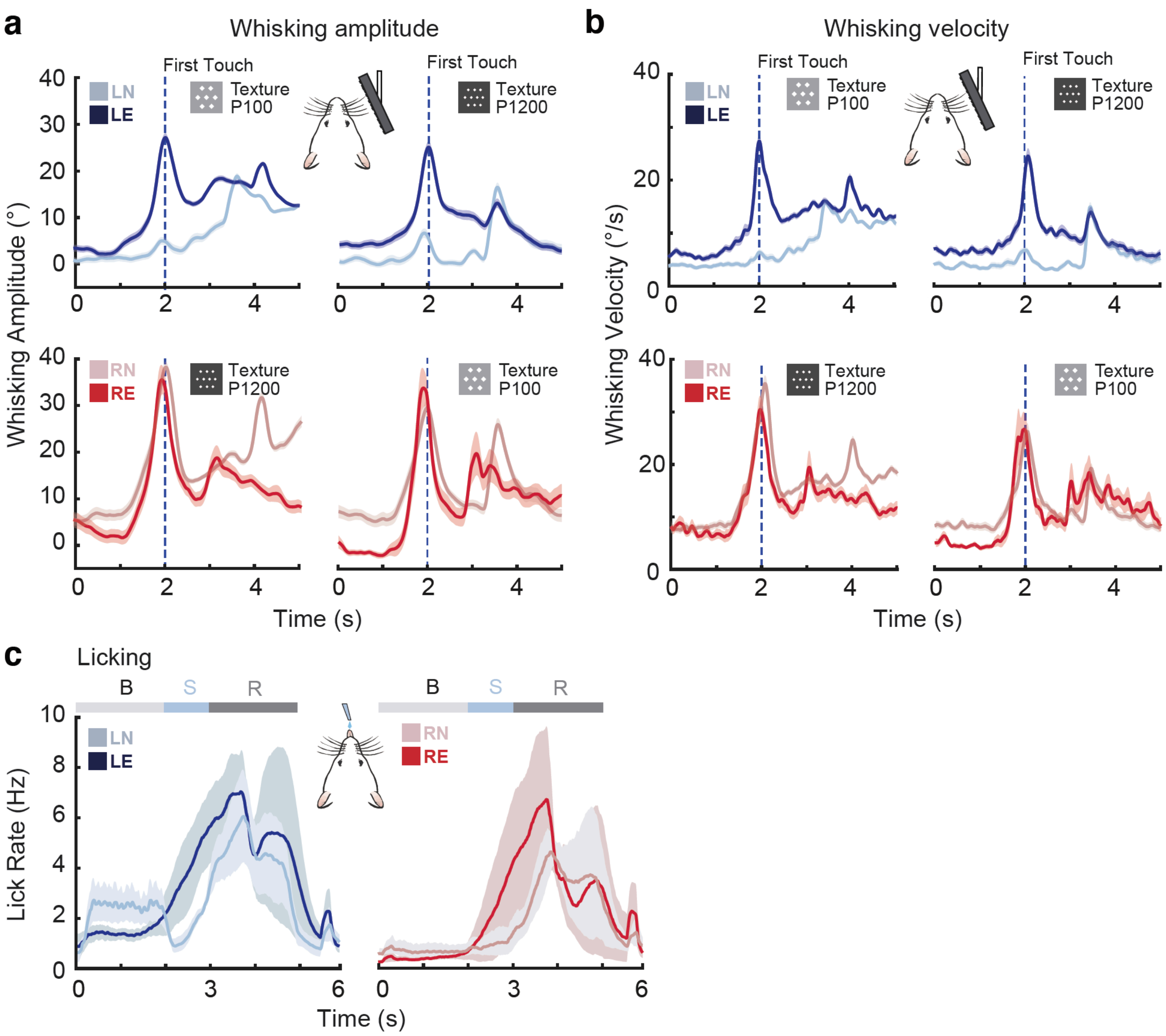


**Supplementary Figure 2 | Whisking and licking behaviour during reversal learning. a,** Upper row: Time-course of envelope whisking amplitude aligned to first-touch during go- (*right*) and no-go-trials (*left*) across two salient periods of initial learning (learning naïve, *LN*; learning expert, *LE*). In naïve animals (*LN*), mice exhibited low amplitude whisking activity throughout most of the trial. In expert mice (*LE*), whisking behaviour became time-locked to the arrival of the texture. Lower row: equivalent whisking traces for the periods after rule switch (reversal naïve, *RN*; reversal expert, *RE*; *right*). Both in *RN* and *RE* periods, mice showed stimulus time-locked whisking amplitude (n = 3 mice). Note that amplitudes and temporal profiles of the whisking envelope were similar for the smooth P1200 and the rough P100 texture, independent of stimulus-outcome association. **b,** Equivalent analysis as in (a) but for the mean whisking velocity. No significant difference was found in the velocity profile between the two textures during the stimulus-presentation period. **c,** Time-course of average lick rates during go-trials across two salient phases of initial learning (*left*) and reversal learning (*right*) (n = 11 mice). Expert mice (*LE* and *RE*) showed both an increase in licking activity during report window (grey) and a decrease of early licks (B-baseline, S-stimulus-presentation, R-reward). Data is presented as mean (solid line) ± S.E.M. (shaded area).

Suppl. Fig. 3

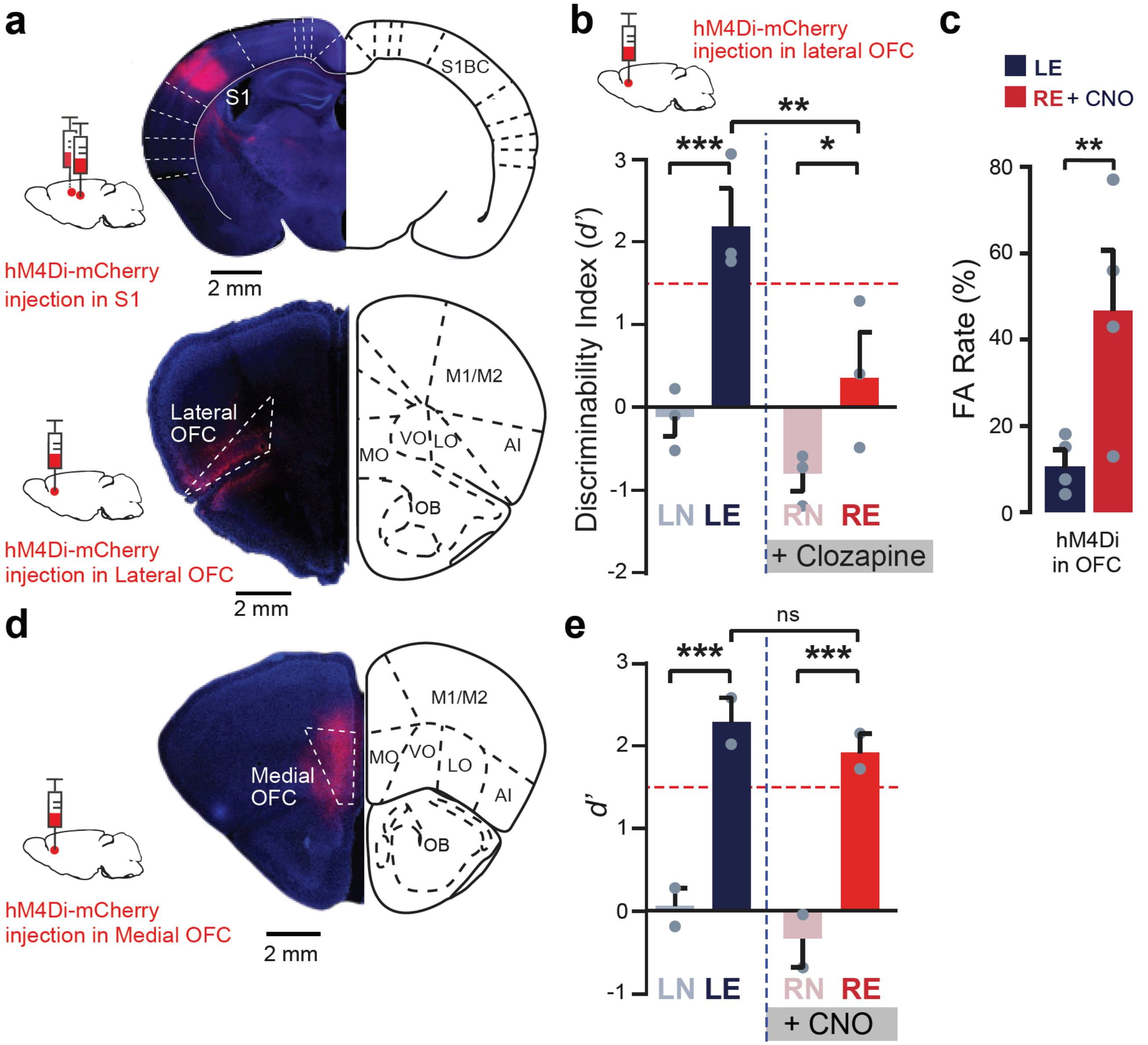


**Supplementary Figure 3 | Immunohistochemical and behavioural validation of pharmacogenetic silencing using inhibitory hM4Di. a,** Neuronal silencing was achieved via viral injection of inhibitory DREADD (AAV-hM4Di-mCherry) into S1 and/or lateral OFC in mice followed by systemic CNO application. S1 injection (top) was bilateral and lateral OFC (LO) injection (below) was unilateral and to the ipsilateral side of the barrel field. **b,** Injection of hM4Di in lateral OFC and systemic administration (i.p.) of clozapine (1-5 mg/kg) after rule-switch (*RN* and *RE*) selectively impaired reversal learning (n = 3 mice). **c,** Injection of hM4Di in lateral OFC and CNO treated animals showed increased perseverative errors (false alarm, FA) in *RE* compared to *LE* (n = 3 mice). **d-e,** Silencing medial OFC (MO) by injecting inhibitory DREADD (AAV-hM4Di-mCherry) unilaterally in the medial OFC followed by daily systemic CNO application after rule-switch in *RN* and *RE* phase, did not have any effect on reversal learning. $^*p < 0.05$, $^{**}p < 0.01$, $^{***}p < 0.001$ Wilcoxon rank-sum test.

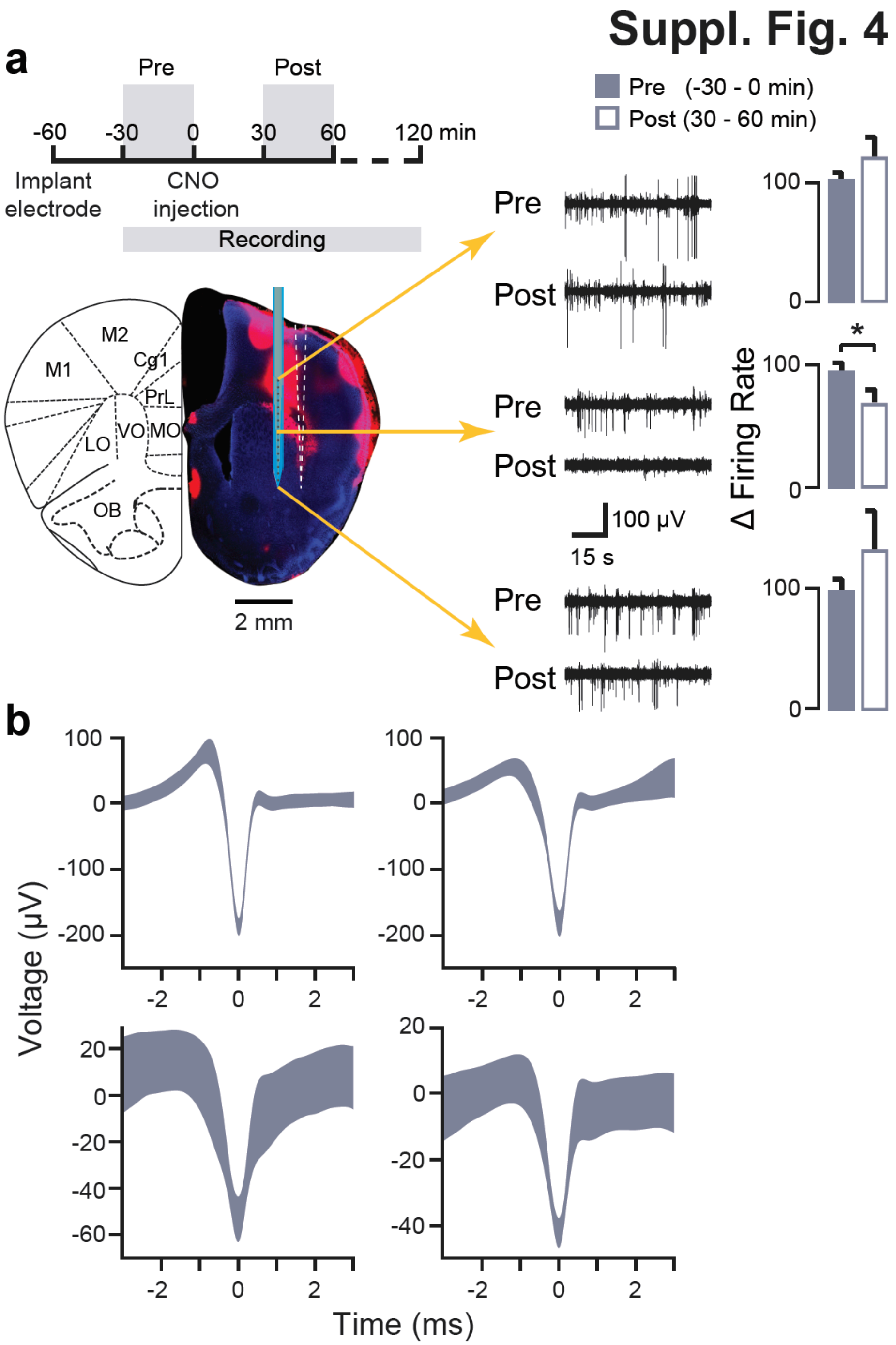


**Supplementary Figure 4 | Electrophysiological validation of lateral OFC silencing using hM4Di. a,** Timeline depicting experimental sequence for validation of lateral OFC (LO) silencing (*top*). Schematic of acute electrophysiological recording from frontal cortex (*bottom*). DAPI stained slice imaged with a confocal microscope showing red fluorescence from DiD to mark the probe location. Example traces from three electrode contacts from one recording session for pre- and post-CNO injection (*middle*). Bar plots showing change in firing rate (% change relative to baseline) for electrode contacts above, in, or below lateral OFC. **b,** Example waveforms from units showing significant modulation by CNO.

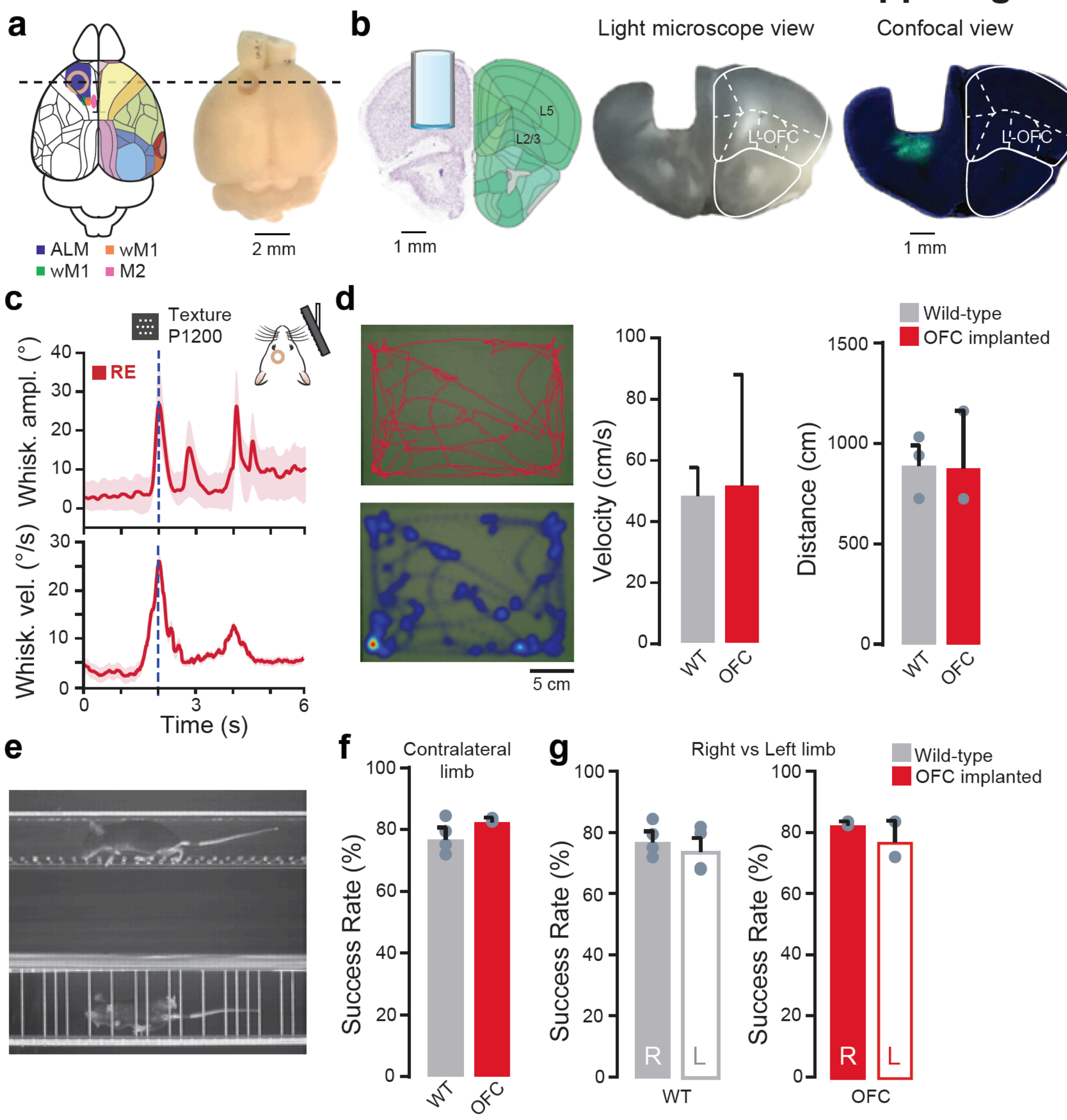


**Supplementary Figure 5 | Unaltered behaviour following OFC cannula implantation. a,** A schematic diagram and whole-brain image showing the location of cannula implantation in OFC. Coloured regions on the schematic indicate pre-motor and motor areas as described in the previous studies[48,49,50,46] (left hemisphere), or regions according to the Allen institute Common coordinate framework (right hemisphere). **b,** A schematic diagram based on the Allen brain atlas, light-microscopic and confocal view shows the GCaMP6 expressing mice in lateral OFC (LO) and cannula placement above the virus injection site. **c,** Whisking behaviour in OFC cannula-implanted animals is preserved. Envelope whisking amplitude (top) and whisking velocity (bottom) in expert animals (*RE*) centred on the texture-approach (n = 2 mice). **d,** Open-field test showed normal locomotor function of wild-type and OFC cannula-implanted mice (n = 4 WT and n = 2 OFC cannula-implanted mice). Representative picture of locomotor track (*top*) and heat-map (*bottom*) of an OFC cannula-implanted mouse. Total distance covered (cm) and mean velocity (cm/s) is plotted. Scale bar = 5 cm. **e,** Horizontal ladder-rung test showed normal locomotor function of wild-type (WT, n = 4) and OFC cannula-implanted mice (n = 2). A representative picture showing paw placement of a mouse on irregular horizontal rung-ladder. **f,** Analysis of paw placement of the limb contralateral to the cannula-implanted side showed no significant difference between wild-type and OFC cannula-implanted mice. **g,** No differences were seen in paw placement of the limb ipsi- or contralateral to the cannula-implanted side in OFC cannula-implanted and in control WT mice. Data is presented as mean ± S.E.M.

**Suppl. Fig. 6**

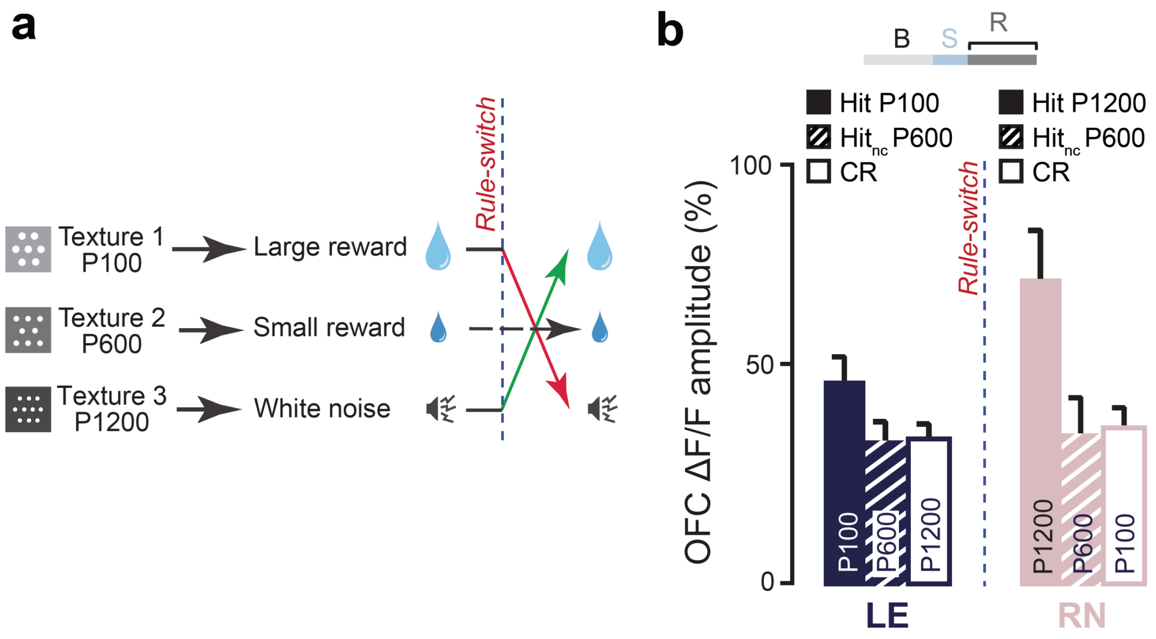


**Supplementary Figure 6 | Re-learning task with neutral context and *in vivo* $Ca^{2+}$ imaging of lateral OFC neurons. a,** Schematic of the stimulus-outcome associations in a three-texture task with positive (large reward) neutral (small reward) and negative (punishment) context. Same coarse P100 and smooth P1200 sandpapers were used as Fig. 1, but an additional intermediate coarse P600 sandpaper was introduced as go-neutral context ($go_{nc}$), whose stimulus-outcome association did not change upon reversal. **b**, Average $Ca^{2+}$ transient amplitude in the reward-outcome window for lateral OFC neurons for Hit, $Hit_{nc}$ and CR trials (n = 63 active neurons out of 228 neurons recorded in three mice; n = 15 sessions) showing increased Hit responses upon rule-switch but no significant changes during $Hit_{nc}$ trials. Across-trial average $Ca^{2+}$ transients for each behavioural period are shown above.

Suppl. Fig. 7

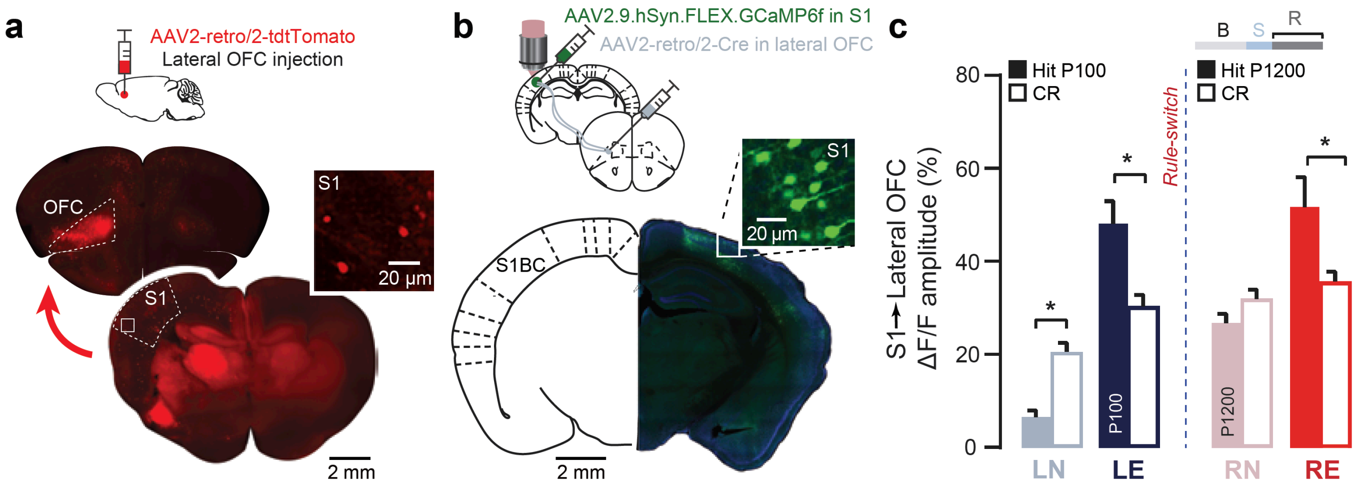


**Supplementary Figure 7 | *In vivo* Ca$^{2+}$ imaging of S1→OFC projecting neurons during reversal learning. a,** Retrograde AAV-retro/2-tdTomato injections *in vivo* in the lateral OFC followed by clearing the brain using CLARITY and whole-brain light-sheet microscopy revealed feed-forward S1→OFC projections from both deeper (L5 and 6) and superficial (L2/3) layers of S1 (n = 2 mice). **b,** S1→OFC projecting neurons were labelled with GCaMP6f using a dual-viral strategy with retrograde AAV2-retro/2-Cre injected in lateral OFC and Cre-dependent AAV-FLEX-GCaMP6f in S1. *Inset*, L2/3 neurons in S1 labelled with such strategy. Average Ca$^{2+}$ transient amplitude in the reward-outcome window shows a sharpening for hit-trials together with a significant increase in amplitude during expert phases of training (*LE* and *RE*), similar to non-selectively labelled S1 neurons (n = 96 responsive neurons over n = 135 recorded in 2 mice, n = 5 sessions/phase). All data presented as mean ± S.E.M., *p < 0.05, **p < 0.01 Wilcoxon rank-sum test.

### Suppl. Fig. 8

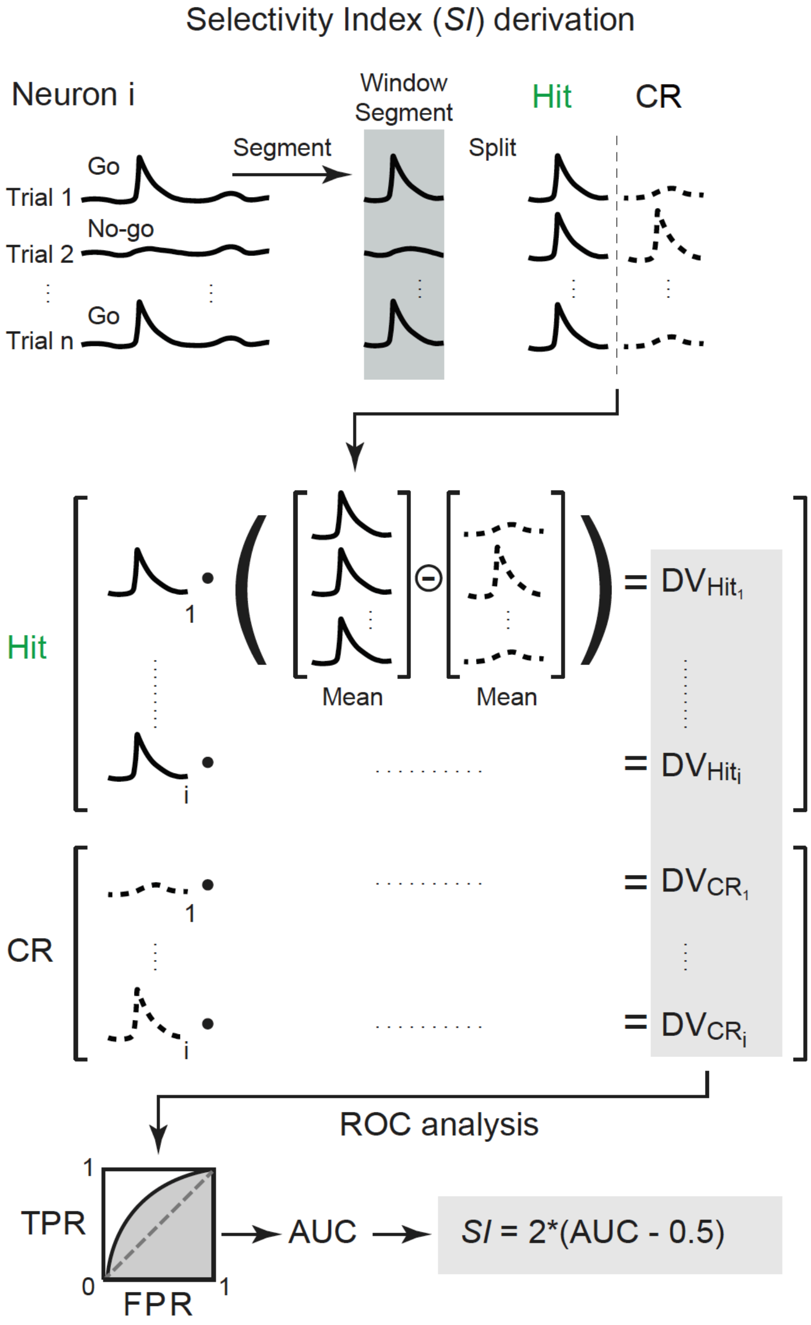


**Supplementary Figure 8 | Schematic of selectivity index calculation.** A schematic view of the step-by-step derivation of the selectivity index (*SI*) from the ROC curves.

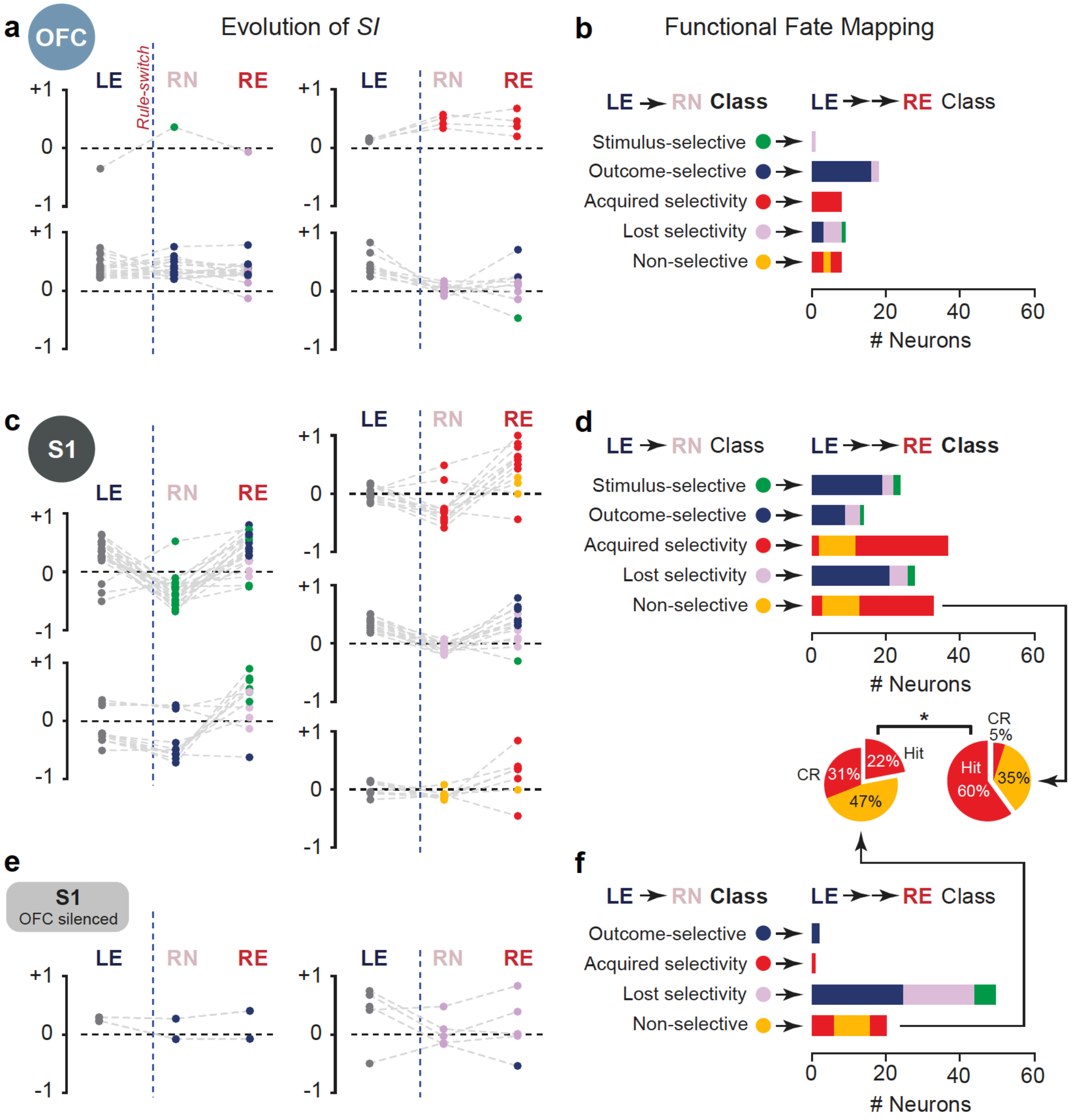


**Supplementary Figure 9 | Tracking neuronal responses during early and late phases of reversal learning. a,** Selectivity indices of longitudinally tracked OFC neurons across the salient task-periods of *LE*, *RN*, and *RE*. Marker colours for *RN* and *RE* indicate the assigned classes for the *LE*→*RN* and *LE*→→*RE* comparisons, respectively. Plots are shown separately for each *LE*→*RN* class. **b**, Fate mapping of longitudinally tracked lateral OFC neurons. For each *LE*→*RN* assigned class, the distribution of these neurons across classes for the *LE*→→*RE* comparison is shown as coloured bar on the right. **c**, Same as in (**a**) but for S1 neurons. **d**, Same as in (**b**) but for S1 neurons. **e**, Same as in (**a**) but for S1 neurons in OFC-silenced mice. **f**, Same as in (**b**) but for S1 neurons in OFC-silenced mice. **Inset in d**, The fate distributions of the non-selective neurons in *LE*→*RN* show a significantly smaller fraction of neurons that acquire selectivity for the newly rewarded go-texture in the *RE* phase in S1 neurons when OFC was silenced in mice (22% vs. 60%, one-tailed Chi-square test). Note that the fate mapping plots include additional neurons compared to (a), (c), and (e) as these were not assigned an *SI* value in each phase but still classified.

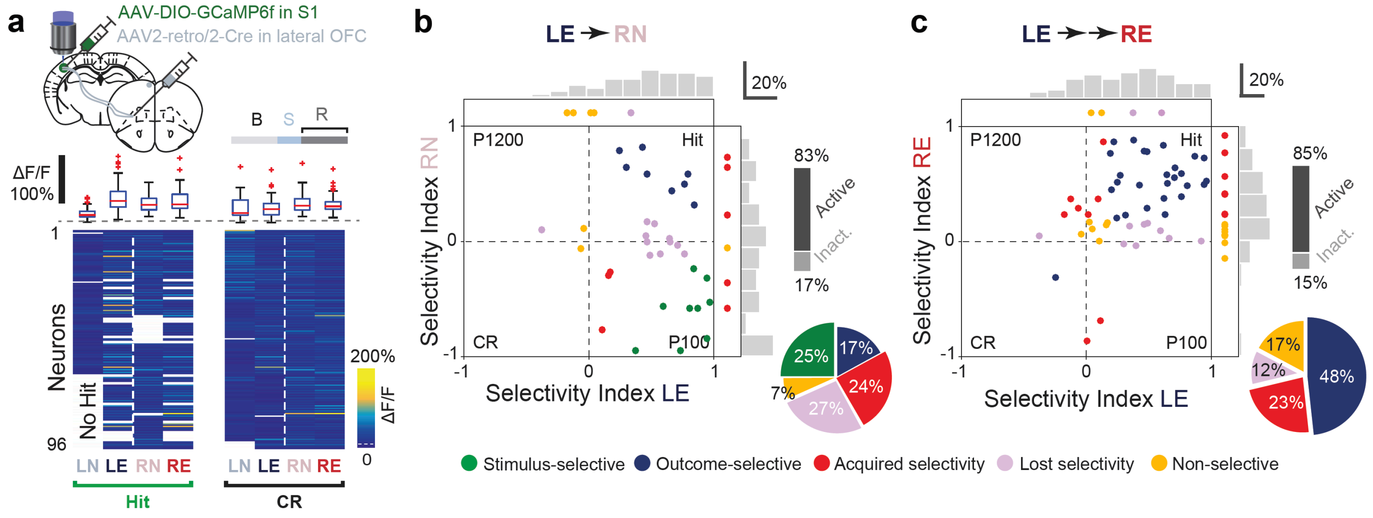


**Supplementary Figure 10 | Task-related functional dynamics in S1→OFC projecting neurons after rule-switch.** **a,** *Top,* S1→OFC projecting neurons were labelled using a dual-viral strategy with retrograde AAV2-retro/2-Cre injected in lateral OFC and Cre-dependent AAV-FLEX-GCaMP6f in S1. *Bottom*, peak responses of S1→OFC projection neurons averaged across hit (left) and CR (right) trials, longitudinally measured across four salient phases (n = 96 neurons from n = 2 mice, n = 5 sessions/phase). Box plots (median, red line; 25$^{th}$ and 75$^{th}$ percentile, box edges; whiskers as most extreme non-outliers; outliers, red crosses; zero, dashed grey line) are also shown (*inset*). **b,** Scatter plot and histogram comparing selectivity index (*SI*) of S1→OFC projecting neurons during learning expert (*LE*) and reversal naïve (*RN*) phase (n = 39 active neurons over n = 46 neurons from n = 2 mice, n = 5 sessions/phase). **c,** Scatter plot and histogram comparing *SI* of S1→OFC projecting neurons during *LE* and reversal expert (*RE*) phase (n = 61 active neurons over n = 73 from n = 2 mice, n = 5 sessions/phase).

Suppl. Fig. 11

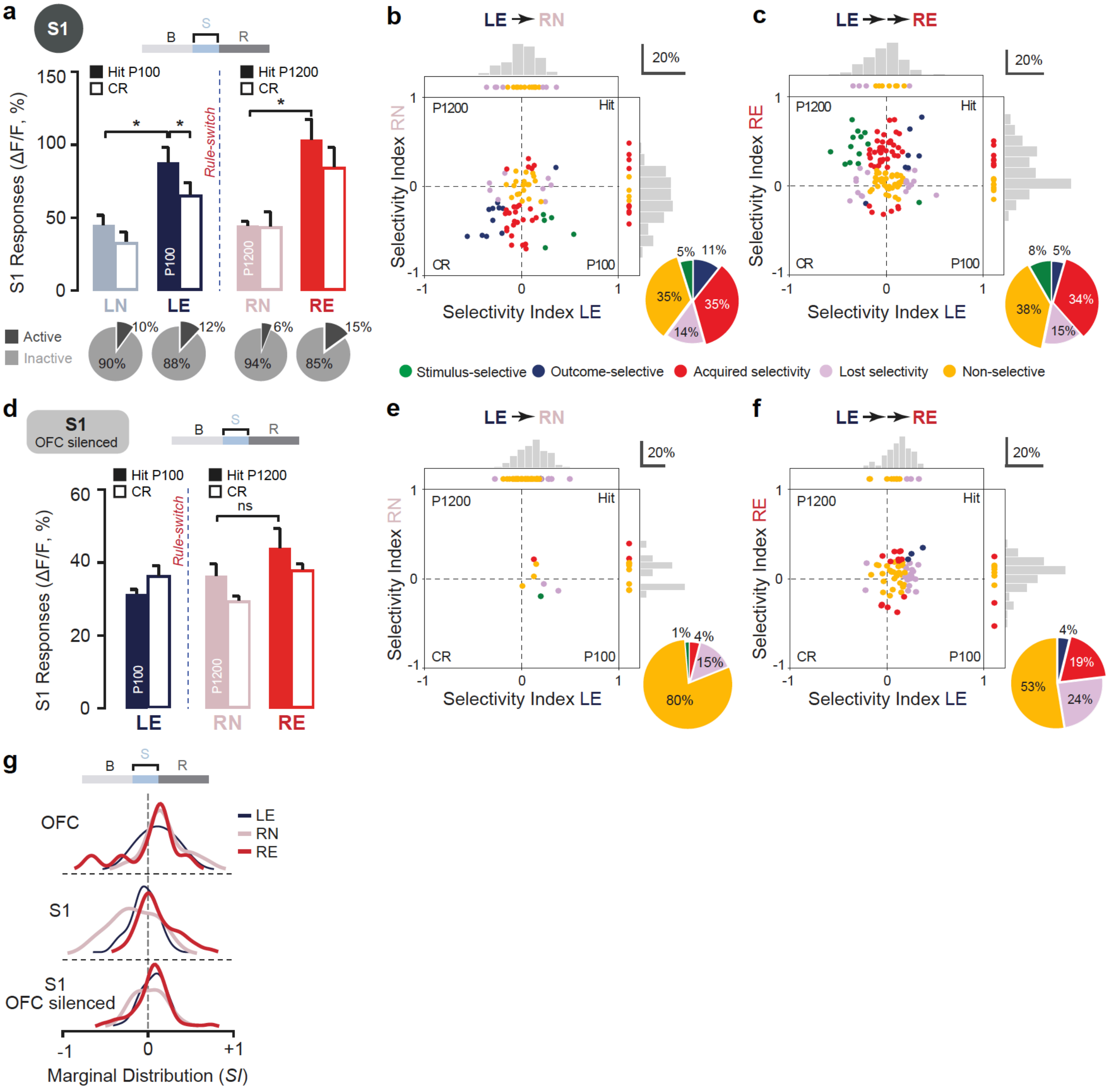


**Supplementary Figure 11 | Texture touch-related dynamics in S1 neurons during reversal learning. a,** Average $Ca^{2+}$ transient amplitude (ΔF/F) in the stimulus-presentation window (S) for S1 neurons (n = 142 neurons in n = 3 mice, n = 2 sessions/phase). **b,** Scatter plot and histogram comparing texture touch-related selectivity index (*SI*) for the stimulus-presentation window for S1 neurons during learning expert (*LE*) and reversal naïve (*RN*) phase (n = 218 from n = 3 mice, n = 28 sessions). **c,** Scatter plot and histogram comparing *SI* of S1 neurons during *LE* and reversal expert (*RE*) phase (n = 218 neurons from n = 3 mice, n = 28 sessions). **d,** Average $Ca^{2+}$ transient amplitude (ΔF/F) in the stimulus-presentation window for S1 neurons in OFC silenced mice (n = 87 neurons in n = 2 mice, n = 2 sessions/phase). **e,** Scatter plot and histogram comparing texture touch-related *SI* of S1 neurons during *LE* and *RN* phase in OFC-silenced mice (n = 165 neurons, n = 25 sessions per phase). **f,** Scatter plot and histogram comparing touch-related *SI* of S1 neurons in lateral OFC-silenced mice during *LE* and *RE* phase (n = 210 neurons in n = 3 mice, n = 28 sessions). **g**, Comparison of *SI* marginal distributions for the three salient periods *LE*, *RN*, and *RE* for OFC neurons (2D scatter plots not shown), S1 neurons (panels c,d), and S1 neurons in lateral OFC-silenced mice (panels e,f).

#### Suppl. Fig. 12

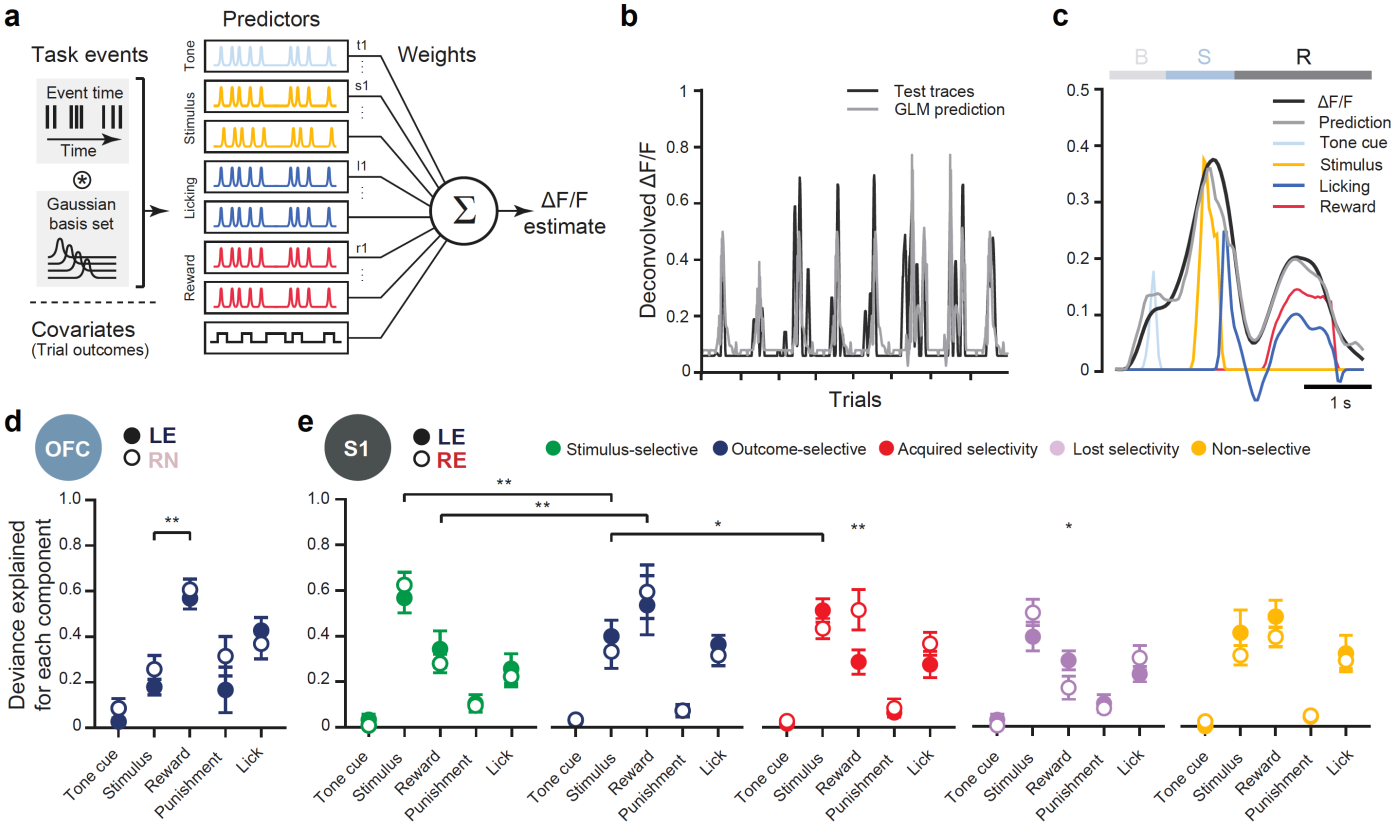


**Supplementary Figure 12 | Differential modulation of task variable-relevant events in neuronal responses. a,** Schematic diagram of a generalised linear model (GLM, Poisson regression) to predict neural activity from behavioural task variables. Each event was expanded into a series of evenly spaced gaussian filters. **b,** GLM predicting deconvolved neural activity of an example S1 outcome-selective (OS) neuron from task variables. **c,** Separate components contributing to the average response of this neuron reveal major sensory modulation together with reward-evoked activity. B-baseline, T-texture touch, R-reward. **d,** To quantify each task variable contribution, the relative fraction of deviance explained is calculated and normalised by the total deviance explained for each neuron both pre- and post-reversal. The reward component in lateral OFC OS neurons is significantly greater than the touch related component. **e,** Fraction of deviance explained for each component in separate subsets of S1 neurons reveal distinct modulations for specific task-related events. Notably, responses of OS S1 neuronal responses are mostly explained by reward component. Licking activity seems to modulate S1 neural responses less than reward in each subset. All data presented as mean ± S.E.M., *$p < 0.05$, **$p < 0.01$, Wilcoxon rank-sum test.

Suppl. Fig. 13

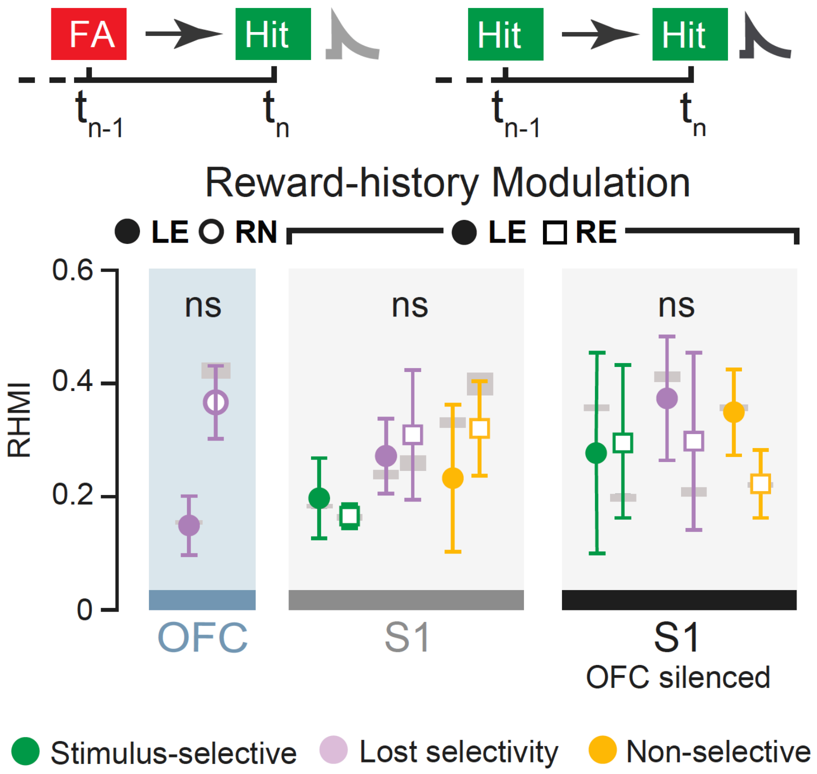


**Supplementary Figure 13 | Reward history-dependent computation in OFC and S1 neurons.** Reward-history modulation index (RHMI) for functional subclasses of neurons in lateral OFC and S1 in OFC intact and OFC-silenced mice ($p < 0.05$; bootstrap-permutation test; S.E.M. of RHMI with permutated indices as grey bars).
